# Scorpio : Enhancing Embeddings to Improve Downstream Analysis of DNA sequences

**DOI:** 10.1101/2024.07.19.604359

**Authors:** Mohammad S. Refahi, Bahrad A. Sokhansanj, Joshua C. Mell, James R. Brown, Hyunwoo Yoo, Gavin Hearne, Gail L. Rosen

## Abstract

Analysis of genomic and metagenomic sequences is inherently more challenging than that of amino acid sequences due to the higher divergence among evolutionarily related nucleotide sequences, variable k-mer and codon usage within and among genomes of diverse species, and poorly understood selective constraints. We introduce Scorpio, a versatile framework designed for nucleotide sequences that employs contrastive learning to improve embeddings. By leveraging pre-trained genomic language models and k-mer frequency embeddings, Scorpio demonstrates competitive performance in diverse applications, including taxonomic and gene classification, antimicrobial resistance (AMR) gene identification, and promoter detection. A key strength of Scorpio is its ability to generalize to novel DNA sequences and taxa, addressing a significant limitation of alignment-based methods. Scorpio has been tested on multiple datasets with DNA sequences of varying lengths (long and short) and shows robust inference capabilities. Additionally, we provide an analysis of the biological information underlying this representation, including correlations between codon adaptation index as a gene expression factor, sequence similarity, and taxonomy, as well as the functional and structural information of genes.

## Introduction

Next-generation sequencing technologies have revolutionized the biological sciences by providing vast pools of genomic and metagenomic data from diverse organisms and environments. Metagenomic data offers the potential to gain insight into the composition and function of microbial communities (“microbiomes”) associated with humans or in the environment. Specifically, shotgun metagenome sequencing from microbial communities, rather than from individual species, enables quantification of in situ microbial consortia to track community diversity, co-evolution, and how community dynamics change in response to environmental perturbations. However, analyzing metagenomic data poses significant challenges. Unlike marker-based community profiling that primarily measures relative abundances of microbial taxa using 16S rRNA amplicon sequencing, shotgun metagenomics provides a more detailed view by capturing the functional genomic content within a community along with associated taxonomic signatures^1^. Alteration in metagenomic content can represent ecological shifts or evolutionary adaptations within species. Handling high-throughput reads, managing the complexity of diverse microbial populations, and resolving genetic differences within taxa are crucial for understanding the functional consequences to changes to the microbiome^2^.

Traditional sequence alignment methods, which align unknown sequences to reference databases of genomic sequence, become computationally difficult with the ever-increasing volume of metagenomic data^3–5^. This motivates the development of alignment-free methods that can rapidly and efficiently characterize sequences found in metagenomic data without relying on computationally expensive alignment processes. Beyond serving as an alternative, such methods can also complement alignment-based approaches by enabling tasks like binning or efficiently identifying key sequences for further detailed analysis^6^. Numerous alignment-free methods have been developed that rely on *k*-mer (i.e., genomic subsequences of length *k*) features. Some of these methods, such as those based on exact *k*-mer matching^2,7^, identify sequences by directly comparing the occurrence of *k*-mers. Others use the composition and abundance of *k*-mers to represent sequences^8^. However, both *k*-mer frequency and exact *k*-mer matching lose positional information—the context and order of *k*-mers within a sequence—which is crucial to the identity and function of genes^9,10^.

To address these limitations, representation learning techniques from natural language processing (NLP) have been adapted for genomic data. By treating nucleotides and amino acids as words in a sentence, models such as Bidirectional Encoder Representations from Transformers (BERT)^11^, Embeddings from Language Models (ELMo)^12^, and Generative Pre-trained Transformer (GPT)^13^ generate lower-dimensional sequence representations through language modeling tasks. These models effectively capture both functional and evolutionary features of sequences but typically require fine-tuning for specific tasks to achieve optimal performance^14–17,17–19^. In recent years, contrastive learning has emerged as a robust technique for refining these representations^20^. This approach involves creating an embedding space where similar sequences are brought closer together, while dissimilar sequences are pushed apart. Contrastive learning enhances the ability to compare sequences rapidly and accurately without relying on traditional alignment methods, especially when implemented using triplet networks,. A triplet network consists of three parallel neural networks that process three inputs: an anchor (a sample from the training set), a positive example (a sample similar to the anchor), and a negative example (a sample dissimilar to the anchor). This structure allows the network to learn to distinguish between similar and dissimilar sequences effectively. By leveraging sequence similarity metrics to optimize the embedding space, contrastive learning with triplet networks has been successfully applied to various tasks in biology, including enzyme activity prediction, identification of disordered protein regions, and protein structural classification^9,21–23^. Another benefit of this approach over other supervised deep learning based models is its generalized and resilient representation, which allows these models to perform well on out-of-domain tasks^24^.

To address challenges of metagenomic analysis, we introduce Scorpio, a flexible framework adaptable to various nucleotide sequence analysis tasks. Scorpio leverages a combination of 6-mer frequency and BigBird embeddings^25^ and is optimized for long sequences. For efficient embedding retrieval, the inference pipeline uses FAISS (Facebook AI Similarity Search)^26^. Scorpio also provides a confidence score for its classifications based on a query-distance and class-probability scoring method, improving prediction accuracy in downstream applications. This framework demonstrates the versatility to analyze both well-characterized sequences and previously unobserved, genetically or taxonomically novel sequences. This capability not only enhances its applicability in metagenomic studies but also reduces the dependency on comprehensive database curation, enabling efficient and accurate insights even in poorly annotated or highly diverse datasets.

We validated Scorpio’s performance on a variety of tasks, including gene identification, taxonomic classification, antimi-crobial resistance (AMR) detection, and promoter region detection. The method proved to be both powerful and efficient when compared to other state-of-the-art methods. Integrating natural language processing techniques with contrastive learning addresses the complex challenges of metagenomic analysis, potentially providing valuable insights into microbial communities and their impact on human and environmental health.

## Results

### Overview of Scorpio

As an initial dataset to evaluate the Scorpio framework, we curated a set of 800,318 sequences. First, we used 1929 bacterial and archaeal genomes curated using the Woltka pipeline^27^, each representing a single genus and with a total of 7.2 million CDS (Figure 1 (a)). Second, gene names alone were used to filter and group protein-coding sequences; unnamed genes with hypothetical or unknown functions were excluded. Third, to improve the dataset’s reliability for training, we included only those genes (497 genes) with >1000 named instances.

**Figure 1.**
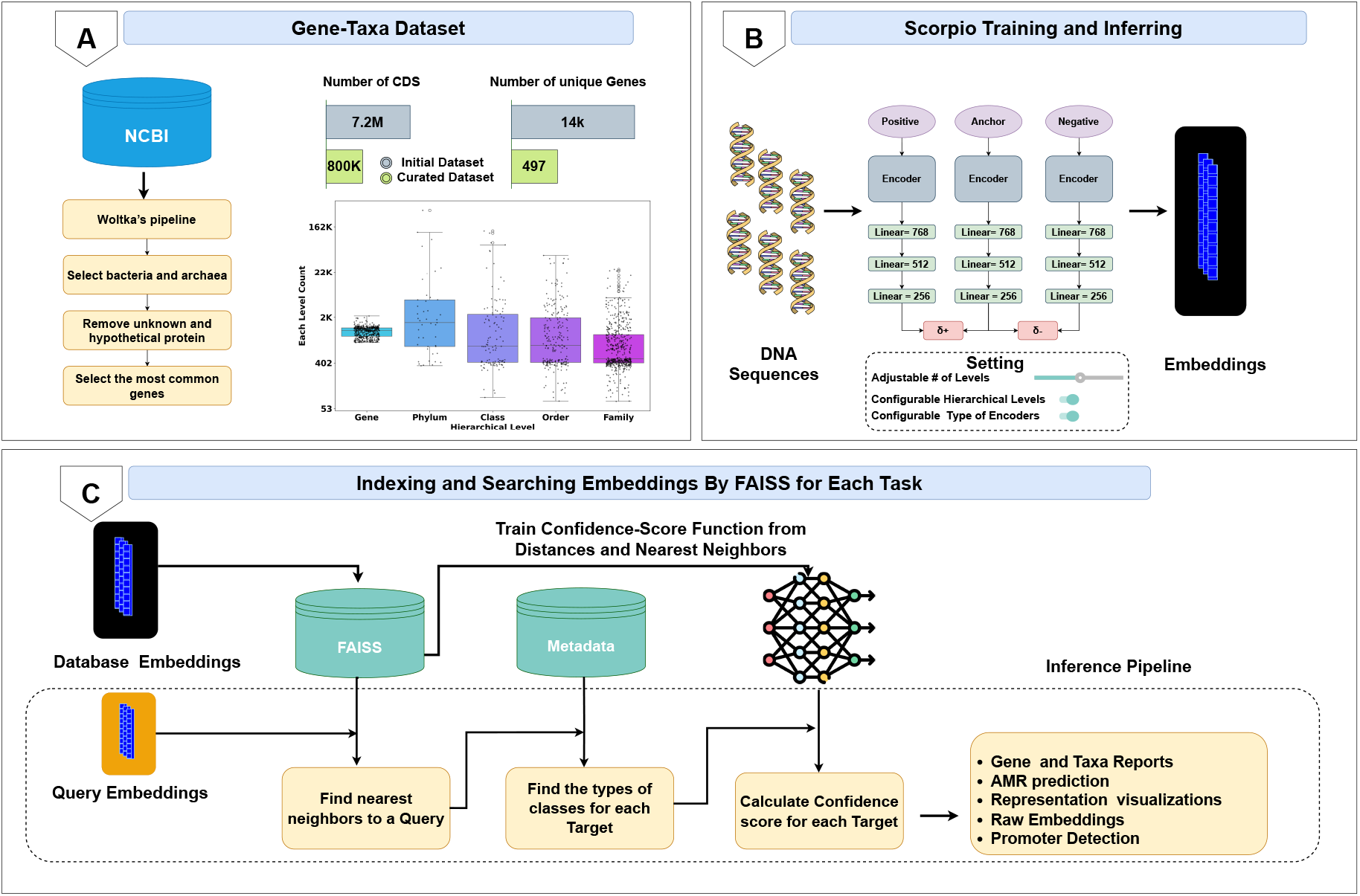
Overview of the Scorpio Framework. (A) Gene-Taxa Dataset Creation: genomes from NCBI was downloaded using the Woltka pipeline27 and filtered to include 497 named genes from 1929 genera (a single-species representative per genus). This process removed most unknown and hypothetical proteins and focused on the most common, conserved, and well-studied genes, particularly housekeeping genes. Genes were labeled with names as-is with no further tests for sequence homology within-label. Results of filtering are shown as a barplot, and the distribution of samples per level is shown in a box plot, indicating a balanced dataset at the gene level. (B) Training and Inferring with Scorpio: DNA sequences are encoded using 6-mer frequency and BigBird embeddings. The configuration supports different Scorpio models, such as Scorpio-6Freq, Scorpio-BigDynamic, and Scorpio-BigEmbed, with adjustable hierarchical levels for enhanced generalization, allowing adaptation to different datasets and hierarchies. During inference, one triplet branch is used to obtain the embedding vector, which is the final layer of the network. (C) Indexing and Searching: FAISS is utilized for efficient embedding retrieval of each query and to find the nearest neighbor. Based on the nearest neighbor from the validation set, we train a confidence score model at each level of the hierarchy. During inference, this model calculates the confidence for each query. Depending on the application,classification results and confidence scores are reported.

**Figure 2.**
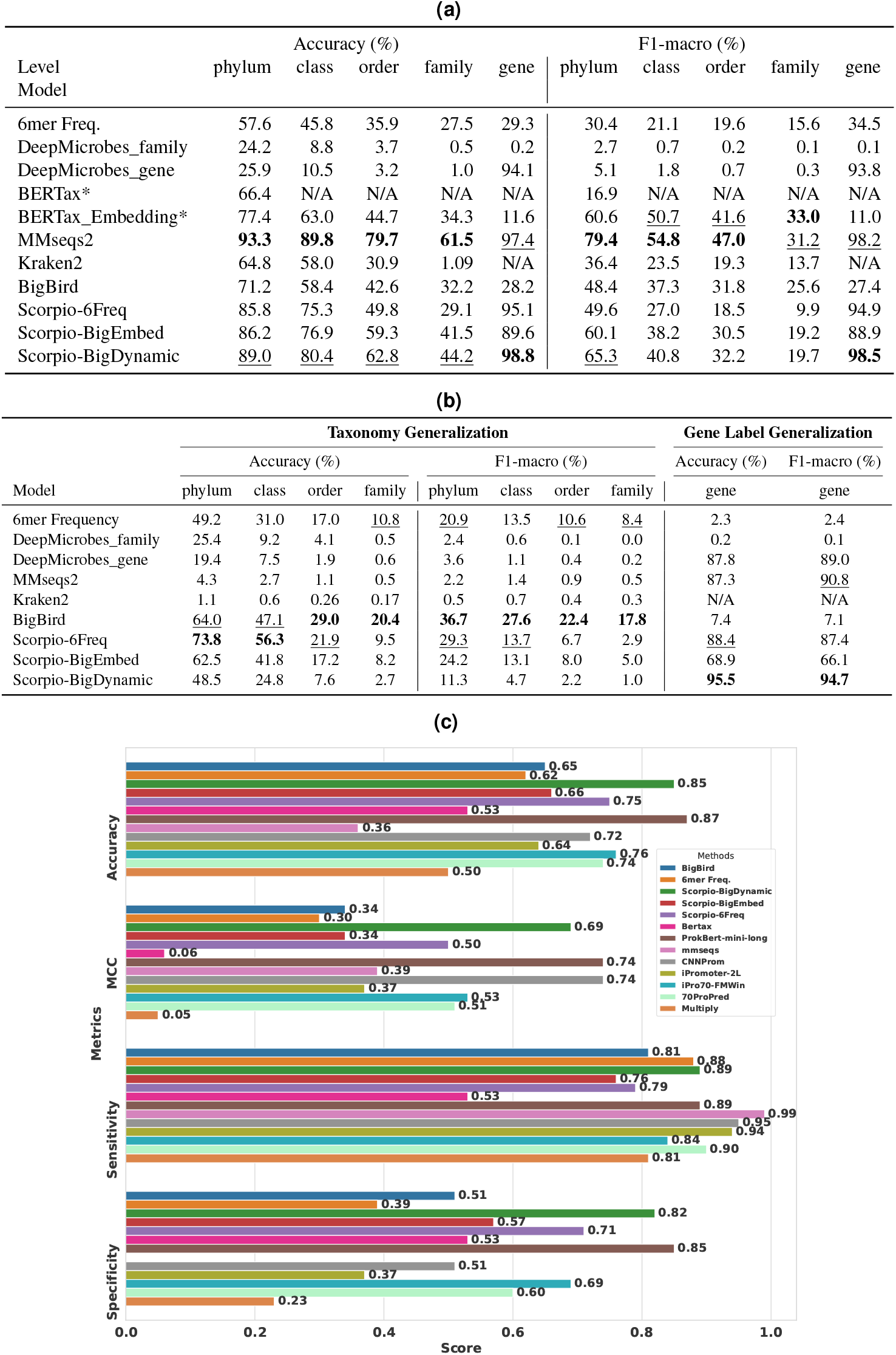
Full gene length results: (a) Memorization Test: Identification of additional training-data-known taxonomy and genes (Test Set). * All models, except for BERTax, were trained on the same dataset; for BERTax, we employed a pre-trained version. (b) Generalization Test: Taxonomy Generalization (Genes-Out Set) and Gene Label Generalization (Taxa-Out Set). We show that while standard techniques, like MMseqs2, memorize data well for identifying known classes, Scorpio is competitive at classifying novel taxa, especially at higher levels and is competitive for genes as well.(c) Performance comparison of different promoter detection methods highlights the effectiveness of our Scorpio approach in handling short-length sequences and out-of-domain tasks for promoter detection

This curated dataset aligns with the study’s goal of addressing both functional and phylogenetic challenges. Phylogenetic biases in some datasets can hinder the ability to recognize and reconcile rare genomes, as shown in various studies such as Centrifuger^28^, fast.genomics^29^, and others^30,31^. These studies highlight how database biases toward specific genomes can significantly impact tool performance. By incorporating fairly responsive genes across taxa, especially those associated with horizontal gene transfer events, we ensure a comprehensive comparison between tools and databases while improving predictions in tasks such as antimicrobial resistance (AMR) prediction^29^. This approach mitigates dataset bias and simulates a scenario akin to few-shot learning, a concept often leveraged in model optimization to enhance performance with limited representative data^32^.

The distribution of instances per class at each level is shown in Figure 1 (a). One advantage of the Scorpio model is the dataset preparation process and the architecture’s ability to train on gene and taxonomic hierarchies. This dual function/taxa focus enables the model to learn multimodal information together, categorized across different hierarchical levels such as phylum, class, order and family,. This preparation of the dataset is the foundation for effectively training the model tocapture the complex relationships in metagenomic data. The training set is carefully balanced at the highest (gene) to lowest (family) level. This balance ensures that we have enough samples for effective triplet training and accurate selection of positive and negative examples.

The framework employs a triplet training approach, where DNA sequences are transformed into embeddings using an encoder mechanism (Figure 1(b)). We have three distinct encoder mechanisms for triplet training: one based on 6-mer frequency (Scorpio-6Freq), and two others based on the embedding layer of BigBird^25^, a transformer architecture optimized for long sequences using sparse attention mechanisms. In one of these BigBird-based mechanisms, we have a fine-tunable embedding layer (Scorpio-BigDynamic), while in the other, all BigBird layers are frozen (Scorpio-BigEmbed). All combinations of positive, anchor, and negative samples are fed into the network to train the triplet network, which processes them through multiple linear layers to fine-tune the embeddings based on the hierarchical labels.

Indexing and searching embeddings efficiently is a critical component of the Scorpio framework (Figure 1(c)). The inference time of deep learning-based methods, particularly those utilizing LLM embeddings, tends to be longer compared to certain conventional bioinformatics tools^2,24^. To address this, we use FAISS (Facebook AI Similarity Search) to store and retrieve precomputed embeddings efficiently. When a query is made, the framework identifies the nearest neighbors to the query embedding and calculates a distance metric. This distance is used to train a simple perceptron model on a range of distance thresholds to predict the F1-macro score. The process normalizes the distance values to confidence scores between 0 and 1, ensuring a robust and interpretable output. The inference pipeline supports diverse outputs, including hierarchical prediction reports and providing raw embeddings for further analysis.

One unique aspect of the Scorpio framework is its flexibility; users can adjust the number of hierarchical levels, select the type of level, and change the order of levels to train the model on. This adaptability is crucial for enhancing the model’s ability to generalize and perform multiple tasks and integrate phylogenetic and functional information effectively. Our evaluations, detailed in the following sections, demonstrate its effectiveness in training for different tasks and its robust adaptability to several potential applications.

### Scorpio embeddings can uncover both the gene’s type and taxonomy levels from full-length gene sequences

Gene-centric metagenomic and pangenomic analysis focuses on identifying coding sequence (CDS) genes from metagenomic datasets or genomic assemblies. With the advancement of long-read sequencing technologies and improved gene-finding algorithms, this approach is gaining popularity and becoming more accessible to researchers.^33–36^. We evaluated the performance of Scorpio embeddings for on a dataset with of the 800,318 full-length DNA gene sequences described above beside these leading methodologies, Kraken2^5^ (a k-mer-based technique widely used for taxonomy), MMseqs2^37^ (a fast and efficient alignment search),DeepMicrobes^10^ (a deep learning technique for taxonomy), and BERTax^19^ (a Transformer-based architecture) (see Methods). For MMseqs2 and all embedding-based methods, we used the best hit for classification. BERTax required a different approach because its original pre-training data did not overlap with our dataset. To fairly evaluate its capabilities, we employed two methods: one leveraging an embedding-based approach integrated with FAISS for best hit classification, and the other utilizing BERTax’s native prediction function to predict taxonomic levels.

In the Test set (Table 2a), we included DNA sequences such that each gene or genus represented was present in the training set, but the specific combinations of genes and genera were not repeated. MMseqs2 had the highest accuracy across taxonomic levels, which was anticipated since alignment-based techniques typically excel with sequences similar to their indexing database. Scorpio outperformed other methods, including Kraken2 and DeepMicrobes. Kraken2’s performance was notably affected by the dataset design, which included only up to one representative gene per genus and only 497 genes. Since Kraken2 relies heavily on large, diverse reference databases with multiple preparations for each taxonomic group, the dataset itself reduced its ability to take advantage of variation across whole genomes.

We next focused on how well the set of methods generalized using the Gene Out and Taxa Out datasets to test performance on previously unseen representative genes and taxa. In generalizing to unknown genes (Table 2b), defined here as novel genes absent from similar genera in the training set, Scorpio embeddings had higher performance than to Kraken2, MMseqs2, and DeepMicrobes, highlighting its ability to capture nuanced patterns within gene sequences, surpassing traditional alignment-based methods that struggle with novel gene classes due to lower sequence similarity with the training set. However, BigBird alone generalized better to classifying taxonomy than Scorpio, having the highest F1-macro across all levels. We attribute this to two main factors: first, the LLM-based embeddings capture more generalized features compared to a strict contrastive learning approach, and second, our model placed gene at the highest level of the hierarchy, so embeddings can become more distinct from each other, reducing performance at the taxonomic levels for out-of-domain data. We also observed this effect when comparing Scorpio-BigEmbed embeddings to Scorpio-BigDynamic, where the latter showed better generalization at lower levels due to hierarchical fine-tuning on top of BigBird, performing better with taxonomy. Notably, these observations also apply to Scorpio-6Freq, which may also be influenced by the hierarchical nature of Scorpio’s training. Scorpio models consistently outperformed others at higher levels of taxonomy, achieving significantly better accuracy performance: 17 times higher than MMseqs2, 67 times higher than Kraken2, and 3 times higher than DeepMicrobes at the phylum level.

In the Taxa Out dataset, which included similar genes but from different phyla than those in the training set„ our Scorpio-BigDynamic model achieved a higher accuracy of 95.5% and an F1-macro score of 94.7%. Interestingly, Scorpio also showed stronger generalization than BigBird in gene classification, with an average performance improvement of 12 times over BigBird. A key advantage of Scorpio over supervised models like DeepMicrobes is its capacity to simultaneously perform taxonomy and gene classification in a single training task by optimizing the loss across all hierarchical gene-taxonomy levels, eliminating the need for separate models for family and gene classification.

### Transferability of Scorpio Embeddings to Other Domains: Antibiotic Resistance Prediction

Next, we evaluated whether Scorpio models originally trained on the **Gene-**Taxa **dataset** in Figure 1 could generate embeddings for antimicrobial resistance prediction tasks. We expected that gene and taxonomy information could help determine the particular genes associated with resistance to a drug class. To test this, we evaluated the transferability of the previously trained gene-taxa model to AMR prediction tasks without additional fine-tuning.

For evaluation, we used a combination of the MEGARes^38^ and CARD^39^ datasets, with details provided in the Dataset section. MEGARes and CARD are global antimicrobial resistance (AMR) databases that integrate relevant data on bacterial taxonomy, genomics, resistance mechanisms, and drug susceptibility. To ensure a fair comparison, we created a custom database by integrating sequences from both MEGARes and CARD, standardizing the data to maintain consistency^40^. We evaluated models based on accuracy and F1-score, choosing benchmark approaches that allowed for database customization. Thus, we evaluated our models with MMseqs2^37^, Abricate^41^ (a well-known tool for mass screening of contigs for antimicrobial resistance genes, virulence factors, and other important genetic markers in microbial genomes), BLASTn^42^ and BERTax^19^.

For embedding-based methods like our BigBird, BERTax, 6-mer Frequency and Scorpios, we used the best hit to determine the class. We present the results in Figure 3a. Scorpio models, particularly Scorpio-BigDynamic and Scorpio-BigEmbed, outperformed all other models in class prediction accuracy across all tasks. While the Scorpio models displayed a 0.4% decrease in F1-macro score of Gene Family Classification, their overall performance, especially in accuracy, consistently surpassed other methods. On average, Scorpio-BigDynamic (gene-taxa) achieved a score of 92.98%, closely followed by Scorpio-BigEmbed (gene-taxa) at 92.83%. In contrast, Abricate exhibited a markedly lower average score of 34.22%, and while Mmseqs2 performed better with an 87.97% average, it still fell short of the accuracy provided by the Scorpio models. An intriguing observation is that LLM-based models (BigBird, BERTax, Scorpio-BigDynamic, and Scorpio-BigEmbed) all outperformed traditional alignment-based tools in classifying resistance mechanisms. This advantage may stem from LLMs’ ability to leverage pre-trained knowledge about patterns associated with resistance that detect functional relationships beyond strict sequence alignment. Notably, resistance genes frequently spread across bacterial species through horizontal gene transfer (HGT)^43^, a process that LLM-based models and Scorpio appear better suited to capture due to their capacity for generalized learning across diverse taxa.

**Figure 3.**
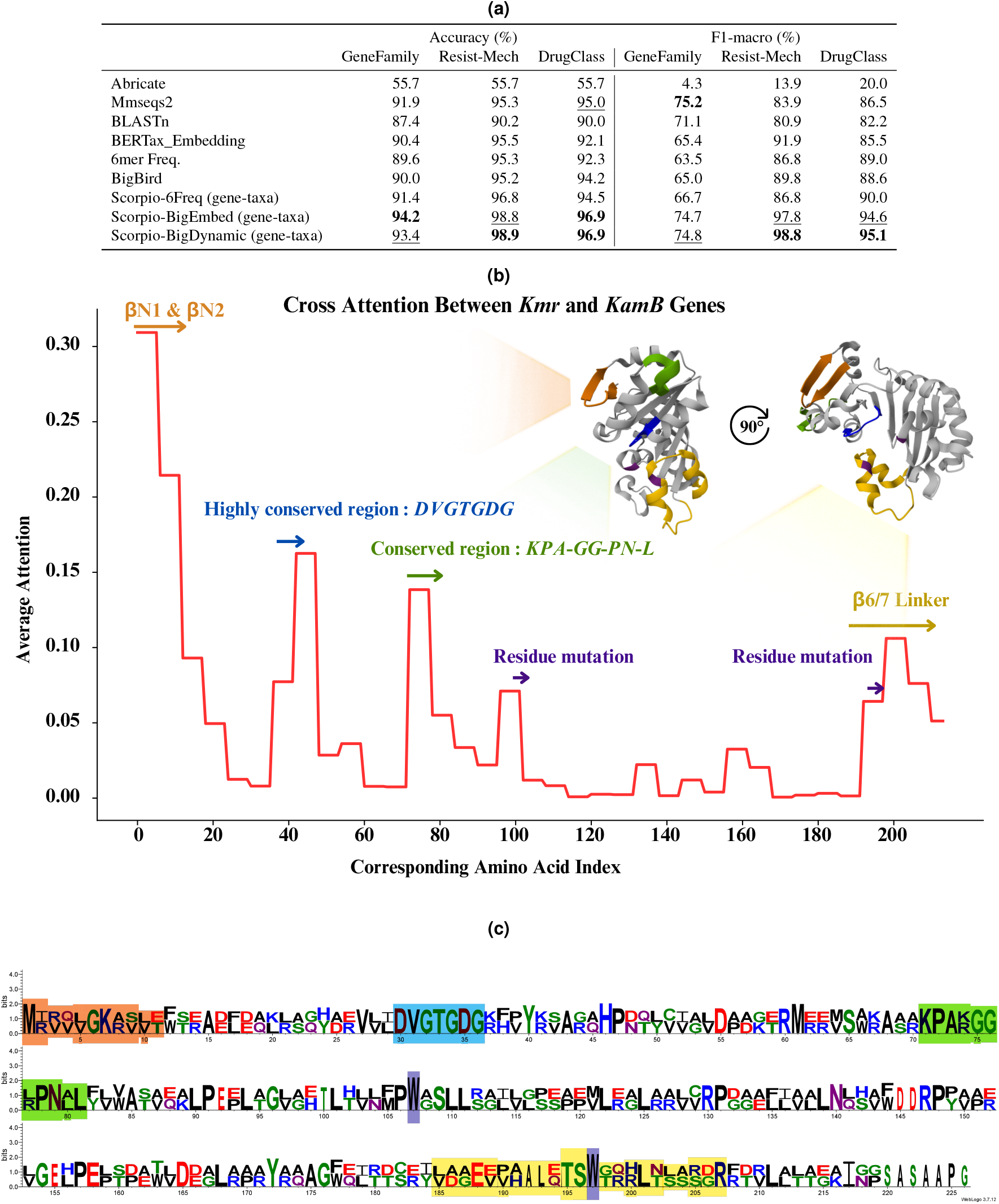
(a) Comparison of Antimicrobial Resistance (AMR) prediction performance metrics across different models. This table highlights that Scorpio models outperform other models in AMR tasks, even though they were not explicitly trained on the AMR dataset, using gene-taxa-based training instead. (b) Cross-attention analysis of two AMR genes (KamB and Kmr) conducted using the Scorpio-BigDynamic model. High-attention regions identified by the model include critical areas such as the *β*^N1^ and *β*^N2^ (orange) regions, conserved regions (blue and green), mutation sites (purple), and the *β*_6/7_ linker (yellow). These regions are predominantly located at the junctions of *α*-helices and *β*-sheets, suggesting functional relevance as detected by the model. (c) Sequence logo of the *KamB* and *Kmr* genes, with letters highlighted based on the regions with high attention identified by the model.

The performance difference between our model and MMseqs2 for resistance mechanism prediction is particularly notable for Antibiotic Target Alteration, with our model achieving a 7% higher accuracy (Supplementary Figure 7). To investigate this further, we analyzed *Kmr* and *KamB*, two AMR genes that share the same resistance mechanism (Antibiotic Target Alteration). *Kmr*, used as a test gene, and *KamB*, included in the training set, were experimentally validated in the study by Savic et al.^44^.

As illustrated in Figure 3c, we identified key regions critical for AMR functionality. These include the *β*_6*/*7_ linker (yellow), which plays an essential role in *S-adenosyl-l-methionine* (SAM) binding and target nucleotide positioning, and the catalytic site at A1408 (purple), a specific nucleotide in the *16S rRNA* that confers resistance to aminoglycosides through methylation^44^. Additional structural features, such as *β* ^*N*1^ and *β* ^*N*2^ (orange), form a *β* -hairpin structure that contributes to protein stability and SAM binding^44^.

Figure 3b highlights the cross-attention analysis^45^ conducted by our Scorpio-BigDynamic model, which effectively captures these regions in two AMR genes (*KamB* and *Kmr*). The analysis uses windowed averages (aggregated for every 6 nucleotides, equivalent to 2 amino acids).Notably, the model demonstrates heightened attention to regions critical for AMR, including *β* ^*N*1^ and *β* ^*N*2^ (orange), as well as conserved regions (blue and green) between the two genes and mutation sites (purple), such as W105A and W193A^44^. These regions are particularly significant due to their conserved or functional importance. We also present the 3D structure of the *KamB* protein in Figure 3b, colorized based on the same high-attention regions identified by our model. Interestingly, these high-attention regions are mostly located at the junctions of *α*-helices and *β* -sheets, suggesting potential functional relevance detected by our model.

In contrast, MMseqs2 failed to predict *Kmr* as a match for *KamB*, likely due to its reliance on strict sequence alignment criteria. The sequence identity after full alignment was only 55%, which falls below the default alignment coverage threshold in MMseqs2 and does not account for differences in codon usage or subtle structural variations. On the other hand, the Scorpio-BigDynamic and Scorpio-BigEmbed models successfully identified *KamB* as the best hit for *Kmr*, showcasing their ability to learn whole-gene representations and effectively capture the structural and functional properties of sequences.

### Fine tuning of Scorpio Embeddings for Bacterial Promoter Prediction

Next, we evaluated Scorpio’s performance in predicting promoter regions that regulate the expression of downstream genes. Notably, since our pre-trained model, BigBird, was originally trained exclusively on gene-encoding regions, it had neither encountered promoter regions nor been trained on sequences of such short lengths prior to fine-tuning. We aimed to investigate the impact of Scorpio on fine-tuning for promoter sequences, considering the significant shift in hierarchical information from coding sequences to promoters. For promoter prediction, the hierarchical representation in Scorpio simplifies to a single level, distinguishing between promoter and non-promoter sequences. We initiated the evaluation by collecting promoter and non-promoter sequences from the ProkBERT dataset^46^. Prokaryotic promoter sequences typically span 81 base pairs. Our model’s performance was independently evaluated using an *Escherichia coli* Sigma70 promoter dataset, providing an objective assessment of its capabilities. This dataset, obtained from the study by Cassiano et al.^47^, comprises 864 *E. coli* sigma70-binding sequences. Positive samples, sourced from Regulon DB^48^, have been experimentally validated and widely recognized promoter sites.

In Figure 2c, we compared the performance results of different methods on promoter detection against our Scorpio method. Firstly, we observe that both Scorpio-BigDynamic and Scorpio-6Freq show improvements in accuracy and Matthews correlation coefficient (MCC) metrics by more than 18% compared to the raw pre-trained model. Additionally, in comparison with state-of-the-art methods^49–53^ for promoter detection, our models perform significantly better on average than most other methods, except for ProkBERT^46^, which had 2% higher accuracy than our model (1% higher sensitivity and 3% higher specificity). This difference likely arises because the BigBird model was solely trained on genes, unlike the ProkBERT model, which was trained on fragments from whole genomes. Language modeling helps capture high-level initial features, and since our model was not exposed to promoter regions during initial training, it may sometimes misclassify promoters as non-promoters. This is further evidenced by the results of Scorpio-BigEmbed, which is trained with frozen embeddings; the model with fixed gene embeddings struggled to adapt to promoter detection. Our pre-trained BigBird model outperforms BERTax, likely due to BERTax’s non-overlapping 3-mer tokenization, which may fail to capture codons in CDS sequences, reducing representation quality and downstream performance. Upon further investigation, we found that some sequences misclassified by our model as non-promoters are actually promoters, such as the sequence, *“CGGTTGCCAACAACGTTCATAACTTTGTTGAGCACCGAT-ACGCATTGTTGGAATTATCGCTCCTGGGCCAGGACCAAGATG”*, which also appears in the coding sequence (CDS) region of *NlpI*. This also suggests that, due to the compact nature of bacterial genomes, embedded promoters may be part of coding sequences. Consequently, a model trained solely on genes might misclassify these sequences since it was not exposed to the broader genomic context during training.

### Scorpio Embeddings Capture Nucleotide-Level Evolution of Coding Sequences and the Relationship Between Codon Adaptation and Sequence Similarity

In molecular evolution, the Codon Adaptation Index (CAI) is an important metric that reflects the frequency of specific codons within a gene in its genomic context and indicates how well-adapted a gene sequence is to its host organism’s translational machinery. Higher CAI values generally correlate with higher RNA expression and more efficient translation^54^. Codon usage bias, the preference for certain synonymous codons, has been shown to regulate transcription and mRNA stability, translational efficiency and accuracy, and co-translational protein folding^55–57^, thus nucleotide-based models capture subtle variations absent from their protein translations.

We thus set out to test whether Scorpio embeddings captured information about CAI following structured approach. First, we selected the 20 most common gene names from our training set. Then, we identified all genera that include these genes, resulting in a dataset covering 31 genera. For each genus, we calculated the CAI of each gene relative to its own genome as the reference^54^. Figure 4a shows the distribution of CAI for each gene across genera sorted by average length (shown as grey bars). This shows that the CAI distribution for these genes is independent of length, which could potentially influence the Scorpio embeddings.

**Figure 4.**
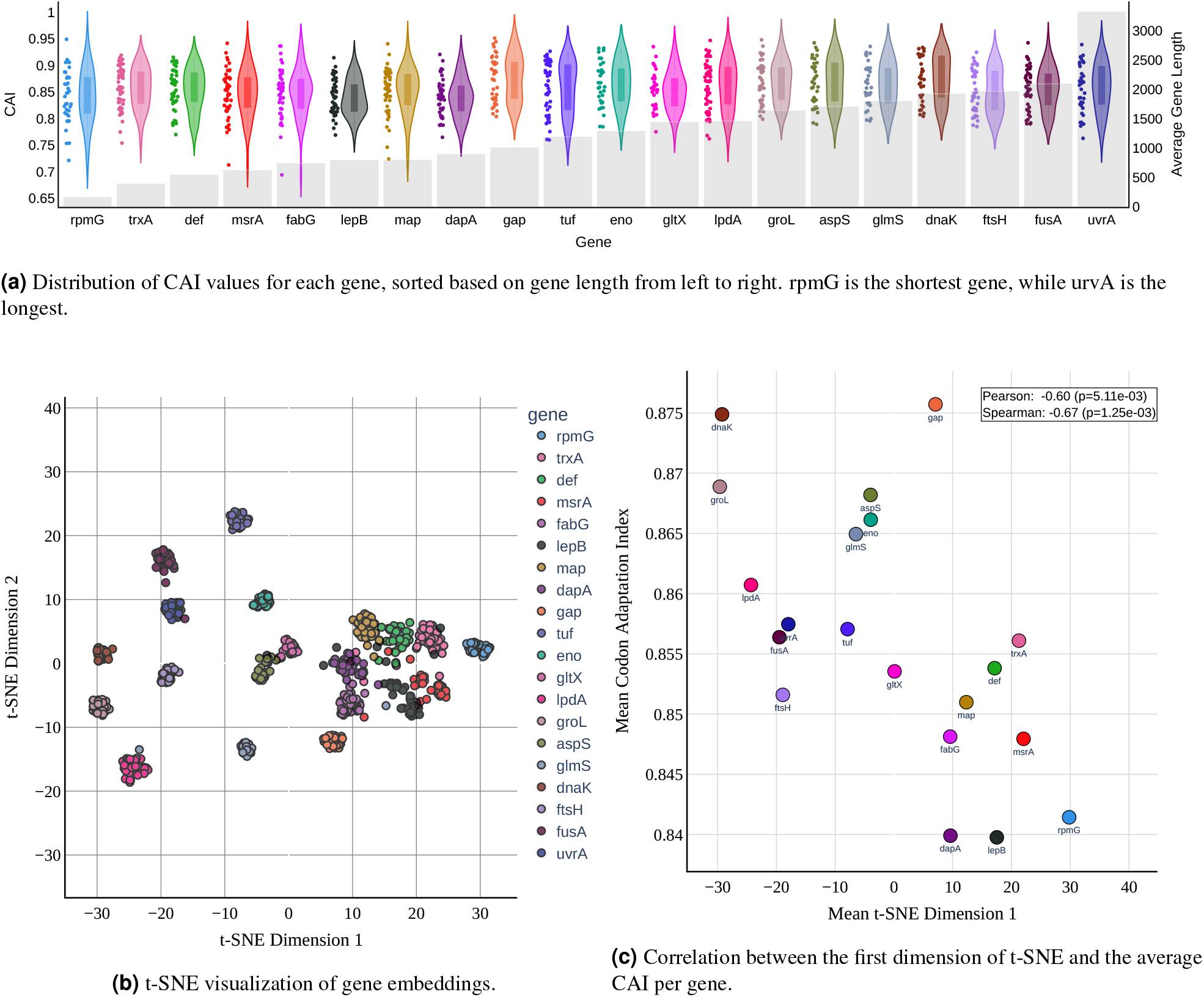
Exploratory analysis shows that Codon Adaptation Index(CAI), independent of gene length metrics, has a significant negative correlation with gene embeddings in the t-SNE visualization, suggesting a potential relationship between gene spatial organization and expression levels. (a) The violin plot shows the distribution of CAI values across genes, indicating variations in codon usage bias. The shaded bars demonstrate that CAI is not dependent on gene length. (b) The t-SNE visualization illustrates gene embeddings in a lower-dimensional space, revealing patterns of similarity and clustering. A high perplexity value was used to capture the global structure of the data, showing how genes relate to each other in space. (c) The correlation analysis between the first dimension of t-SNE embeddings and CAI values provides insights into the relationship between gene spatial organization and CAI. This analysis suggests a significant correlation between gene expression levels and CAI, with Pearson and Spearman’s rank correlation coefficients of -0.60 (*p* = 5.11×10^−3^) and -0.67 (*p* = 1.25×10^−3^), respectively.

Next, to obtain a global representation of the sequences, we employed t-SNE. Although t-SNE transformations of embeddings (Figure 4b) can be nonlinear and may depend on parameters like perplexity and the number of iterations, we used a perplexity value of approximately 50 to capture global structural information in our embedding space^58,59^. A higher perplexity value helps obtain more global rather than local information about each cluster and their distribution in the space. In Figure 4c, we compared the average CAI against the average of the first t-SNE dimension of our embeddings. A negative correlation emerged, with Pearson and Spearman’s rank correlation coefficients of -0.60 (*p* = 5.11*e*^−3^) and -0.67 (*p* = 1.25*e*^−3^), respectively. This indicates a significant negative correlation between the overall representation of these 20 genes’ embeddings in space and the CAI.

Although we observed a correlation, it should be emphasized that embeddings do not linearly represent sequences. Understanding the causal influences for the observed gene distributions in the embedding space is highly complex and involves multiple factors. For example, in Figure 4b, we noticed that AspS and GltX genes, both encoding aminoacyl-tRNA synthetases-key enzymes in the translation of the genetic code-are clustered closely together in embedding space despite not having similar CAI. Additionally, we observed distinct clustering when considering a substantial portion of the aminoacyl-tRNA synthetases, along with others likely involved in tRNA biosynthesis. Other proteins, including several ribosomal proteins, also appeared near each other in the embedding space (Supplementary Figure 9), suggesting that the embeddings capture structural and functional information about genes.

Our analysis also indicated a correlation between the embedding distance of genes and their sequence similarities. We used edit distance to measure gene distance and Euclidean distance to measure embedding distance, as shown in (Supplementary Figure 10). Our examination of sequence similarity within each gene shows that genes in embedding space are distributed based on their sequence similarity. On average, the *R*^2^ value is about 0.41, indicating a moderately large correlation between the edit distance of gene pairs and the Euclidean distance of their embeddings.

These analyses suggest that factors such as gene expression, function, taxonomy information, and sequence similarity may influence the organization of genes in the embedding space. However, it is important to caution that the relationship between sequences and their embeddings is not a simple one-to-one mapping, and the non-linear transformation complicates direct interpretations.

### Assessing Confidence Scores: A Comparative Analysis of Gene and Taxonomy Classification Methods

For benchmarking against existing algorithms and improving useability on metagenomic datasets, we introduce a novel confidence scoring method for classifications based on Scorpio embeddings^60^. Since the gene-level class was trained with only 497 gene labels, evaluating the quality of classifications is crucial for profiling metagenomic reads that come from all genes across all genomes in a community. Most methods like Kraken2 and MMseqs2 apply a cutoff threshold using a confidence score of E-value before reporting results. Using such a threshold presents a trade-off between the number of sequences classified and the precision but is especially required in the presence of many off-target sequences.

We evaluated the gene and taxonomy identification performance of our method against established approaches Kraken2 and MMseqs2. For Kraken2, our training set comprising 540K sequences was indexed, setting the confidence parameters as minimum-hit-groups 1 and confidence as 0. With MMseqs2, we employed the easy-search method on same indexed dataset, specifying search-type 3 for nucleotide/nucleotide searches, while retaining default values for other parameters. We adjusted our threshold for confidence reporting to observe differences in the number of classified sequences and precision (see Methods) We show the results in Figure 5e, displaying the number of classified sequences for each method. Both Kraken2 and MMseqs2 encountered challenges in classifying genes from the Gene Out dataset, with Kraken2 detecting only 2.8% and MMseqs2 detecting 28%. This underscores the drawbacks of methodological factors such as sequence similarity and long k-mer searching in accurately classifying novel sequences. In Table 5a, we show the precision of our model compared to others at different threshold values. Even though the number of classified sequences in the Gene Out dataset is very low, the precision for both Kraken2 and MMseqs2 is also low compared to our model across various thresholds. Scorpio achieved 94% precision and 90% precision when we used thresholds that returned the same number of classified sequences as Kraken2 and MMseqs2. For the Taxa Out dataset, which consists of sequences from similar genes but from unseen taxa, we observe that MMseqs2 was highly effective, correctly classifying 99% of sequences, while Kraken2 classifies only 50%. Considering precision, which crucially reflects the accuracy of classifications without being influenced by the dataset’s total size, MMseqs2 is particularly strong at identifying similar genes. Scorpio also performs well, though with lower precision than MMseq2 for this task and dataset. Kraken2 classified more sequences than Scorpio, since it shares similar phyla and genes with the training set, but Scorpio still outperformed Kraken2 in precision, even when returning a similar number of sequences. Unlike other methods, our approach selects thresholds by considering both the number of sequences to return and the precision. By calculating a confidence score specific to the dataset, we determine a cut-off based on test sets. This threshold represents an inflection point, balancing the number of classified sequences while maintaining high precision. Alternatively, all results can be returned with the associated confidence score and user-defined cut-offs.

**Figure 5.**
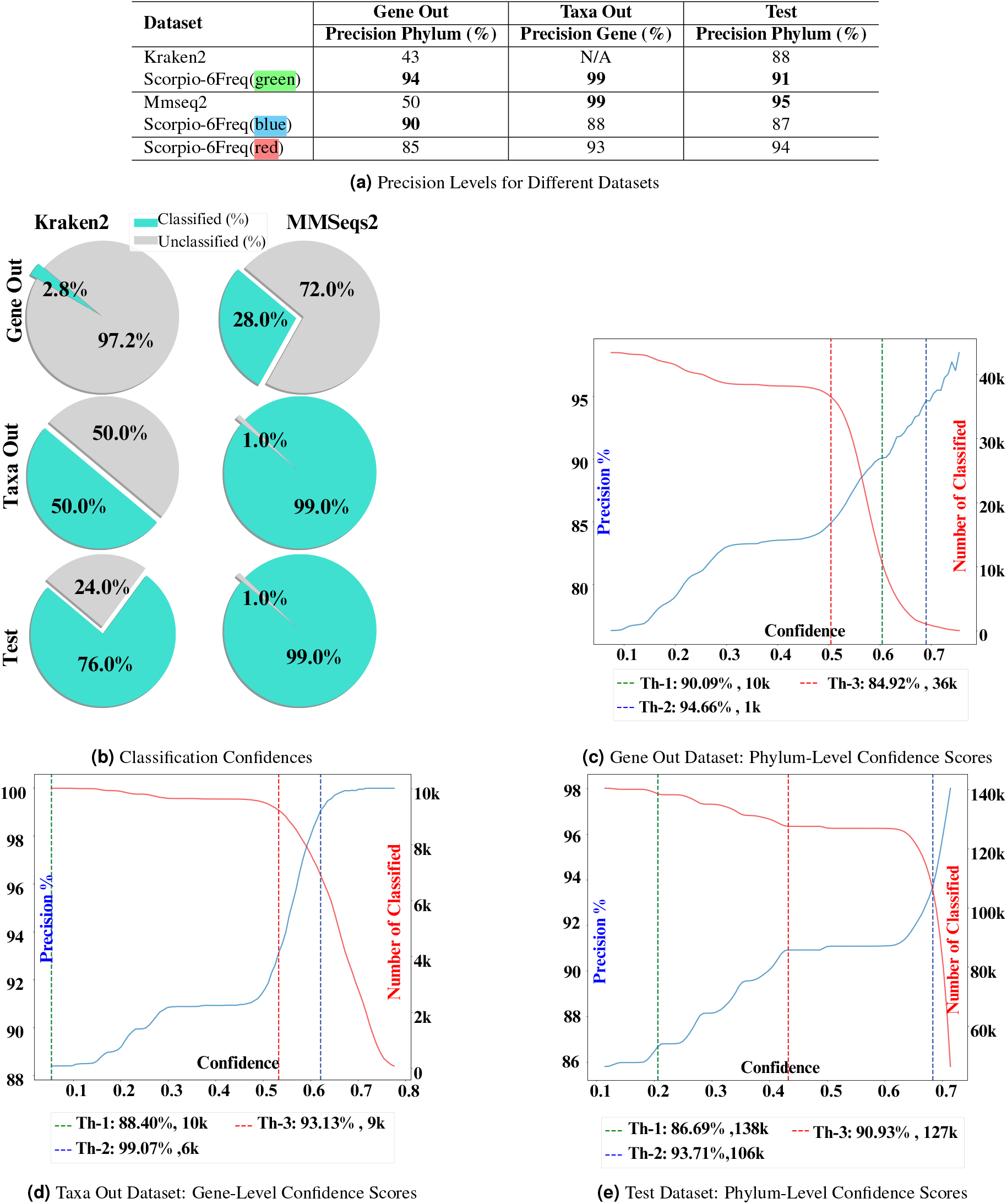
We examined Kraken2 and MMseq2 thresholds and their impact on the number of classified instances. As shown in subpanel (b), the number of classified instances varies across datasets for each method. In the “Gene Out” dataset, Kraken2 classified 2.8% of instances, while MMseq2 classified 28%. We compared our model’s precision across these different thresholds (subpanels c, d, and e), with green representing Kraken2-like thresholds, blue for MMseq2-like thresholds, and red for our thresholds. In Table (a), at a 2.8% classification rate (green), our Scorpio model achieved 94% precision, compared to Kraken2’s 43%. At a 28% classification rate (blue), our Scorpio model achieved 90% precision, while MMseq2 achieved 50%. This analysis demonstrates our model’s effectiveness in maintaining high precision while balancing the number of classified instances in novel sequences.

In Figure 5, we show precision vs. confidence and the number of classified sequences vs. confidence for all three datasets. We also highlight the thresholds s used based on how many sequences were classified by Kraken2 and MMseqs2. In all plots, the red threshold indicates the selection based on MMseqs2, and the blue threshold indicates the cutoff based on Kraken2. The red is the threshold we pick based on the infimum between the number of classified sequences and precision. As it is observable in Figures 5c, 5d, and 5e, all three cases exhibit an increasing trend between the confidence score and precision, validating the reliability of the score.

### Scorpio embeddings can identify both gene and taxonomy labels of short fragments

We next evaluated the effectiveness of Scorpio embeddings in identifying genes and classifying taxonomy in short fragments to test the potential of using Scorpio for critical tasks in metagenomics. We focus on a target read size of 400 bp, comparable to overlapping paired reads generated by Illumina and other short-read as some next-generation sequencing (NGS) platforms. All models used for testing are trained and indexed on the same dataset, except for BERTax:, where we used the pre-trained BERTax. Detailed information about the dataset and the choice of 400 bp read length can be found in the dataset section.

One significant advantage of our approach revealed in by this analysis is the reduced training time compared to DeepMicrobes. As illustrated in (Supplementary Figure 8), the unified objective across all hierarchical levels in Scorpio eliminates the need for separate model training for each task, which is necessary for DeepMicrobes. This streamlined process enhances both efficiency and scalability, making Scorpio a powerful tool in metagenomic analysis.

For the test set (Table 1a), MMseq2 outperformed other methods at various taxonomic levels. Scorpio-BigDynamic excelled at the gene level and was second-best at the phylum level. The 6-mer Frequency representation, despite not using a learning procedure, performed well, indicating its effectiveness for memorization with similar training-testing sequences. Kraken2 showed high precision at lower taxonomic levels. However, Scorpio’s hierarchical selection was significantly affected by dataset imbalance at lower taxonomic levels. Some studies suggest that using balanced datasets is essential for training contrastive learning-based models^22^.

**Table 1.** Short fragment length (400bp) results: (a) Memorization Test: Identifying additional examples of training-data-known taxonomy and genes (Test Set); **\*** All models, except for BERTax, were trained on the same dataset; for BERTax, we employed a pre-trained version. (b) Generalization Test: Taxonomy Generalization (Gene Out Set) and Gene Label Generalization (Taxa Out Set) Tests. Again, Scorpio is superior at classifying novel organisms at the phylum level and beats out every method for the gene level.

| Level<br>Model | Accuracy (%) |  |  |  |  | F1-macro (%) |  |  |  |  |
| --- | --- | --- | --- | --- | --- | --- | --- | --- | --- | --- |
|  | phylum | class | order | family | gene | phylum | class | order | family | gene |
| 6mer Freq. | 90.3 | 86.1 | 76.7 | 65.6 | 92.4 | 78.3 | 69.0 | 63.4 | 55.8 | 91.9 |
| BigBird | 72.4 | 62.7 | 50.1 | 41.5 | 55.6 | 53.6 | 44.0 | 40.1 | 35.5 | 54.9 |
| DeepMicrobes_family | 72.7 | 61.6 | 43.6 | 30.8 | 3.1 | 42.7 | 28.3 | 24.2 | 19.7 | 2.4 |
| DeepMicrobes_gene | 21.5 | 8.9 | 2.4 | 0.7 | 93.0 | 4.3 | 1.4 | 0.5 | 0.2 | 93.2 |
| BERTax* | 76.4 | N/A | N/A | N/A | N/A | 22.9 | N/A | N/A | N/A | N/A |
| BERTax_Embedding* | 55.2 | 38.4 | 20.9 | 13.0 | 15.5 | 27.9 | 16.8 | 12.0 | 8.4 | 14.8 |
| Mmseqs2 | **94.8** | **92.3** | **84.1** | **70.7** | <u>97.7</u> | **85.0** | **75.8** | **69.4** | 57.9 | <u>97.2</u> |
| Kraken2 | 77.8 | 74.4 | 66.7 | 59.6 | N/A | 70.0 | 67.2 | <u>64.4</u> | **60.9** | N/A |
| Scorpio-BigEmbed | 76.6 | 66.1 | 49.8 | 38.9 | 74.0 | 54.3 | 42.4 | 37.1 | 31.4 | 74.5 |
| Scorpio-6Freq | 81.3 | 70.4 | 47.7 | 32.6 | 92.2 | 49.8 | 34.6 | 29.1 | 22.9 | 92.3 |
| Scorpio-BigDynamic | <u>91.0</u> | 83.4 | 63.3 | 45.8 | **98.8** | 73.7 | 53.3 | 42.9 | 32.8 | **98.9** |

**Table 1.** Short fragment length (400bp) results: (a) Memorization Test: Identifying additional examples of training-data-known taxonomy and genes (Test Set); \* All models, except for BERTax, were trained on the same dataset; for BERTax, we employed a pre-trained version.
| Model | Taxonomic Generalization |  |  |  |  |  |  |  | Gene Generalization |  |
| --- | --- | --- | --- | --- | --- | --- | --- | --- | --- | --- |
|  | Accuracy (%) |  |  |  | F1-macro (%) |  |  |  | Accuracy (%) | F1-macro (%) |
|  | phylum | class | order | family | phylum | class | order | family | gene | gene |
| 6mer Freq. | 47.0 | 29.9 | 14.9 | 8.4 | 11.4 | 8.1 | 6.8 | 5.1 | 54.7 | 52.7 |
| BigBird | 51.4 | 33.9 | 17.8 | <u>10.6</u> | 14.3 | <u>11.7</u> | 9.8 | <u>7.5</u> | 16.5 | 14.4 |
| DeepMicrobes_family | <u>55.8</u> | <u>40.3</u> | **22.1** | **13.2** | <u>15.7</u> | **11.8** | **10.3** | **8.3** | 2.8 | 1.9 |
| DeepMicrobes_gene | 14.2 | 5.9 | 1.8 | 0.5 | 2.1 | 0.9 | 0.4 | 0.1 | 76.0 | 77.0 |
| Mmseqs2 | 2.6 | 1.8 | 0.8 | 0.4 | 1.0 | 0.6 | 0.5 | 0.3 | <u>78.5</u> | <u>86.1</u> |
| Kraken2 | 0.93 | 0.63 | 0.27 | 0.11 | 0.2 | 0.1 | 0.09 | 0.06 | N/A | N/A |
| Scorpio-BigEmbed | 54.8 | 37.2 | 18.0 | 9.6 | 14.5 | 10.0 | 7.4 | 5.2 | 41.7 | 42.2 |
| Scorpio-6Freq | 50.0 | 31.1 | 9.4 | 4.0 | 10.7 | 6.1 | 3.1 | 1.5 | 72.4 | 73.5 |
| Scorpio-BigDynamic | **61.2** | **43.1** | <u>18.3</u> | 7.8 | **17.8** | 10.4 | 5.6 | 2.7 | **92.3** | **93.1** |

The test set illustrates different methods’ memorization capabilities, while the Genes-Out and Taxa Out datasets demonstrate generalizability. In the Gene Out set (Table 1a), Scorpio outperformed others at the phylum and class levels in both accuracy and F1 score. DeepMicrobes_family excelled at lower levels like order and family, as it is specifically trained for the family level, unlike our model, which is trained on six levels of hierarchy. DeepMicrobes_gene, trained for the gene level, showed low performance across all taxonomic levels. Our model significantly outperformed MMseq2 and Kraken2, with over 60% improvement at the phylum level. This improvement is due to MMseq2 and Kraken2 struggling with novel gene sequences not in their databases, whereas Scorpio’s embeddings, which capture k-mer frequency and sequence similarity, performed much better than sequence search methods. For the TaxaOut set (Table 1), our Scorpio-BigDynamic model outperformed other models in gene classification, achieving 92% accuracy, while MMseqs2 and DeepMicrobes_gene models achieved 78% and 76% accuracy, respectively. These tests show the generalization of these algorithms on more challenging previously unseen sequences.

With Scorpio embeddings, we also gained valuable interpretability insights into our model’s ability to discriminate both genes and the taxonomy of short fragments. In Figures 6a and 6b, we compare the t-SNE representations of embeddings from both BigBird and Scorpio-BigEmbed models. In Figure 6a, distinct clusters of genes are clearly visible, a distinction that is not as apparent in the pre-trained model. This highlights the robust nature of our model in differentiating gene sequences. Additionally, when we colorize the visualization based on phyla, as seen in Figure 6b, it becomes evident that Scorpio is adept at detecting taxonomic information compared to the pre-trained model. Although there are slight differences between embeddings of each taxonomic level, Scorpio’s hierarchical structure enhances its ability to distinguish taxa within each gene cluster. This hierarchical clustering is particularly effective, demonstrating that our model performs significantly better in taxonomy-based differentiation compared to the pre-trained model.

**Figure 6.**
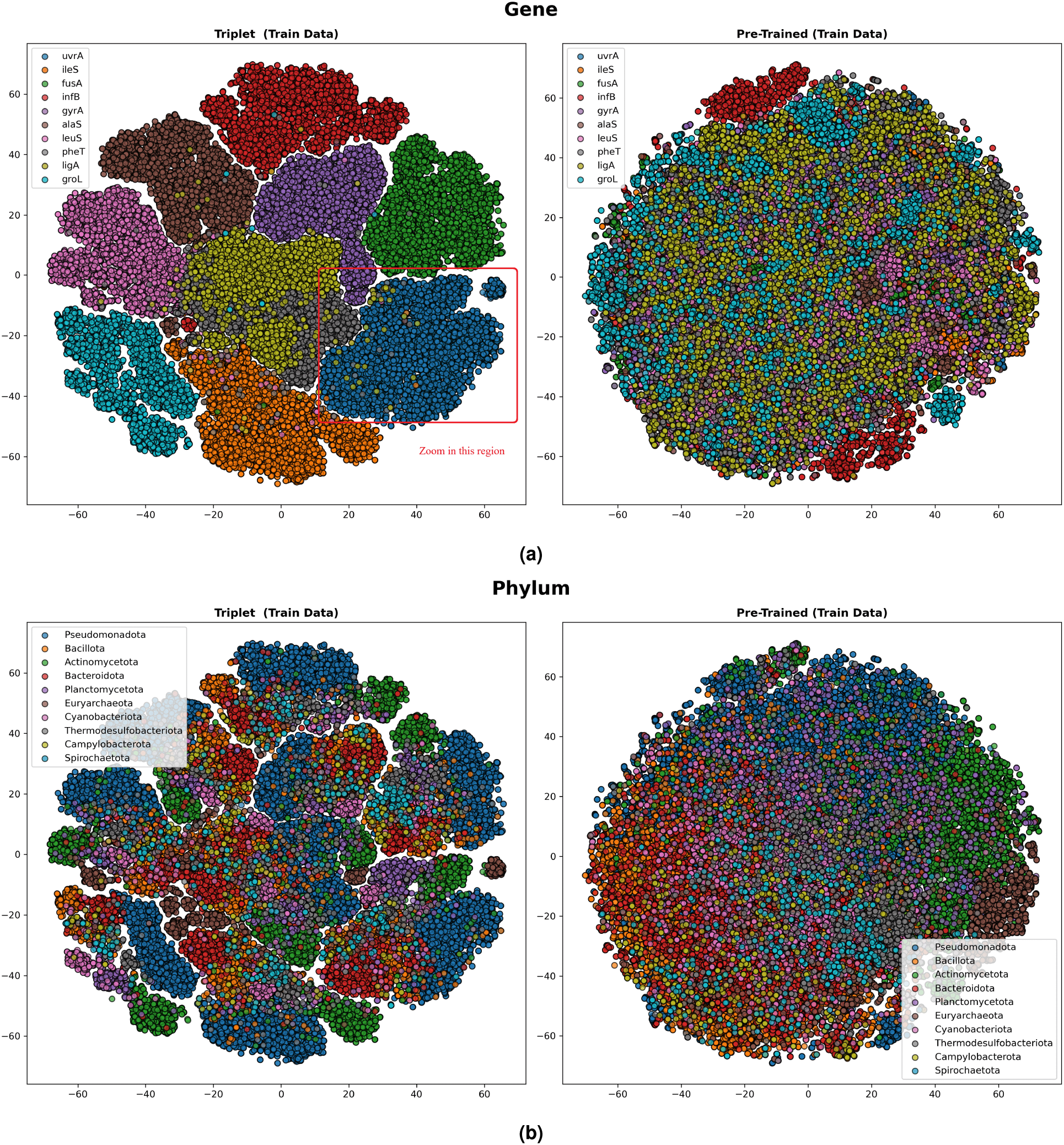
(a)t-SNE visualization of embeddings colorized based on the 10 most common phyla in the dataset using the Scorpio-BigEmbed(Triplet) and BigBird(Pre-Trained) Models(b) t-SNE visualization ofembeddings colorized based on the 10 most common gene in the dataset using the Scorpio-BigEmbed(Triplet) and BigBird(Pre-Trained) Models

As a proof of concept to support future experiments, we conducted an experiment using ART-simulated data^61^ with 150 bp reads. Details of the dataset and results are provided in Supplementary Table 3. Our results demonstrate that Scorpio generally achieves higher accuracy across all taxonomic levels compared to Kraken2. For instance, Scorpio achieved a phylum-level prediction accuracy of 27.9%, significantly outperforming Kraken2’s 0.15%, which classified only approximately 0.38% of the sequences. Despite these promising results, Scorpio’s lower F1-macro scores may reflect its sensitivity to sequencing errors, sequence length dependencies, and challenges in embedding-based search, particularly when handling long-read embeddings for short-read searches in underrepresented classes.

One intriguing insight from this visualization, using the 10 most common phyla, is that in Figure 6b, Euryarchaeota displays a highly distinct representation compared to Bacteria. Despite the model’s attempt to cluster the genes, the hierarchical clustering ability and the significant distinction between Archaea and Bacteria prevented the model from clustering sequences from the same genes but different Kingdoms together. This indicates that taxonomic information influences the model’s organization. To further explore these insights, we zoomed in on a region in Figure 7 that includes all fragments of the *urvA* gene. We colorized this region based on different hierarchical levels, each time displaying the 10 most common categories for each level. It is noteworthy how the model discriminates based on each category at lower hierarchical levels. However, it becomes apparent that as we delve into lower hierarchy levels, the sample size for each genus becomes limited, reflecting the initial gene-based discrimination.

**Figure 7.**
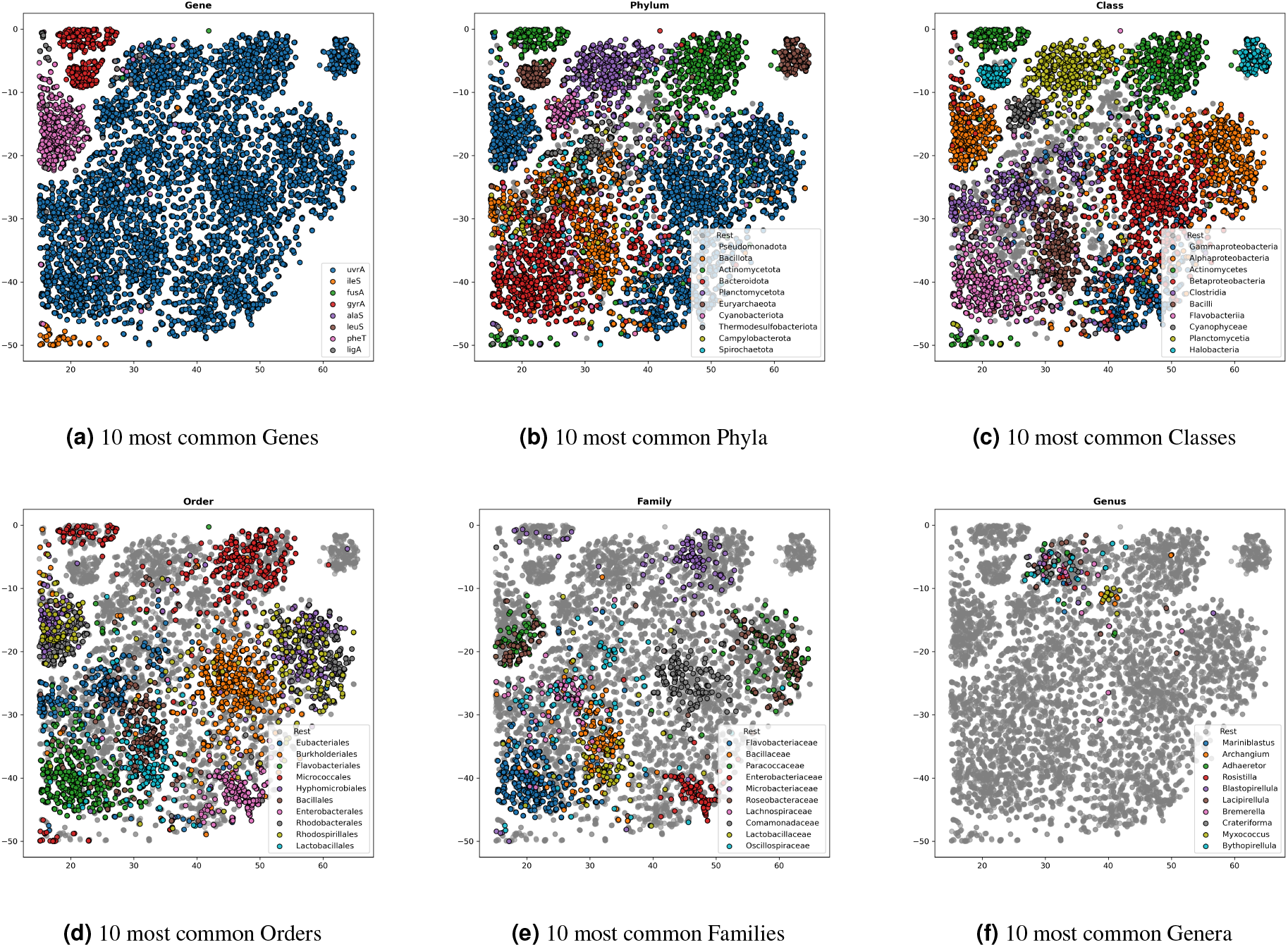
Analyzing a region which includes the *urvA* gene in the t-SNE plot of Figure 6a and colorizing for each level.

## Discussion

In this study, we introduced the Scorpio framework, which leverages exiting pre-trained language models with triplet networks and contrastive learning to enhance the analysis of DNA sequence data. The adoption of nucleotide-based models, unlike traditional protein-focused models^62,63^, could contribute to our understanding by capturing nuances associated with gene expression and translational efficiency embedded with the the nucleotide sequences that encode proteins. By using both pre-trained language models and k-mer frequency embeddings, we aimed to demonstrate the robustness of our framework across multiple types of encoders, all of which showed promising results. We showed that Scorpio, despite only one training round on protein-coding gene sequences with both gene and taxa labels, is among the top-performing algorithms across a variety of tasks. Specifically, Scorpio significantly improved taxonomic and gene classification accuracy, particularly in out-of-domain datasets, thereby showcasing the robustness and generalizability of the method. The superior performance of Scorpio at gene identification and competitive performance for taxonomic classification can be attributed to the ability of triplet networks to learn more discriminative features by optimizing the distance between positive and negative pairs. This capability is crucial when dealing with the high-dimensional and complex nature of metagenomic data, where traditional alignment-based methods may fall short.

One of the key strengths of the Scorpio framework is its versatility in handling multiple tasks. By extending the triplet network to specific applications such as antimicrobial resistance prediction and promoter detection, we demonstrated the adaptability of our model to various different nucleotide sequence analysis tasks. Our model generalization surpasses methods like MMseqs2, Kraken2 and Abricate, which rely on k-mer and sequence similarity searches, by showcasing the effectiveness of embedding-based search.

The ability to transfer learning from one domain to another and fine-tune the model for specific tasks highlights the potential of the Scorpio approach for practical applications in health and environmental diagnostics. This adaptability could be particularly valuable in metagenomics, where rapid identification and characterization of novel genes and taxa are critical. Our evaluation across diverse datasets with varying gene lengths demonstrates the robustness of our method. Consistent performance improvements across different datasets indicate that our model can effectively generalize from training data to unseen data, a crucial requirement for reliable metagenomic analysis.

Additionally, our framework incorporates a novel confidence scoring mechanism, which provides a measure of the reliability of the results. This scoring method uses both the distance to the nearest neighbor and the class probabilities derived from neighboring embeddings, ensuring that confidence scores are meaningful and reliable for use as a quality estimator of the search method.

There are several areas for future research and development. Firstly, while our model performed well in out-of-domain classification tasks, further improvements could be achieved by increasing the dataset size to include a more curated and diverse genes-taxa. Incorporating additional sources of biological information, such as functional annotations and protein interaction networks, could enhance the interpretability of the learned embeddings and provide deeper insights into the functional roles of genes and taxa. Furthermore, expanding the data for underrepresented classes and employing techniques to address data imbalance could improve accuracy at lower levels of the hierarchy. Secondly, the computational efficiency of our framework could be optimized further. Although the use of FAISS for efficient embedding retrieval was effective, and the possibility of running FAISS on GPU makes it faster, exploring more advanced indexing techniques like ScaNN (Scalable Nearest Neighbors)^64^, which has demonstrated promising results in our comparative analysis with FAISS in the Supplementary Materials, or parallel processing strategies, could reduce the computational overhead and enable the analysis of even larger datasets. Lastly, while our approach has shown significant promise, it should be extended and fine-tuned to other domains beyond gene/taxa/AMR/promoter classification. The principles underlying our Scorpio framework could be applied to other types of biological sequence data, such as transcriptomics and proteomics, potentially opening up new avenues for research and application. We also plan to test Scorpio on experimental samples and make our tools available as practical applications for integration into metagenomics pipelines.

In conclusion, our study presents a robust and versatile framework for DNA sequence classification, leveraging triplet networks with contrastive learning and integrated embeddings from PreTrained language models and k-mer frequencies. This approach significantly advances our capacity to process and interpret complex microbiome data, offering valuable insights for health and environmental diagnostics. Future work will focus on further optimizing the model, integrating additional biological information, and exploring its applicability to other domains of genomic research.

## Method

### Scorpio: Architecture

The architectural design features three layers in each branch. It maps either a 768-dimensional pre-trained embedding or a 4096-dimensional 6-mer frequency vector to a 256-dimensional Scorpio embedding. Each encoder block consists of a linear layer followed by a ReLU activation function, producing 256-dimensional embedding vectors for downstream analyses. The architecture is flexible and can easily handle k-mer frequency data with only small changes. Specifically, by altering the size of the first layer, the model can handle different input dimensions. For example, with 6-mers, the first layer is 4096-dimensional, while the rest of the architecture remains consistent. This flexibility demonstrates the model’s ability to adapt to various data representations, as illustrated in Figure 8a. We employ three types of encoders. The first is the 6-mer frequency encoder, which calculates the 6-mer frequency(Scorpio-6Freq)representation and passes it through the architecture. The other two are variations of the BigBird model: one with a trainable final embedding layer(Scorpio-BigDynamic), and another with all layers frozen(Scorpio-BigEmbed). Both BigBird models use an average pooling layer to aggregate embeddings into a single vector, maintaining the model’s simplicity and efficiency.

### Scorpio: Triplet Training

In the realm of training contrastive learning models, the method for choosing triplet sets (anchor, positive, negative) is crucial for shaping the model’s ability to understand semantic connections. Inspired by the Heinzinger et al.^22^, our approach involves a refined selection process with adjustments to enhance versatility across different similarity levels.

In our training set, each sample serves as an anchor during every epoch. We deliberately randomize the selection of positive and negative samples for each anchor in each epoch, ensuring the model encounters new instances while revisiting the same triplet set. This repetition guards against overfitting and promotes generalization. Our hierarchical sample selection involves a two-step process depicted in Figure 8b. First, we randomly select the similarity level for each anchor. Based on this similarity level, we then randomly select positive and negative samples. Positive samples come from the same hierarchy level as the anchor, while negative samples are selected from one level higher than the anchor’s similarity level. Importantly, similarity at a lower hierarchical level does not necessarily guarantee similarity at higher levels. For example, choosing sequences from the same phylum for positive and negative samples implies similarity at the gene level as well. This careful consideration ensures that positive samples not only share specific traits but also align across all hierarchical levels, reinforcing a nuanced understanding of the hierarchy. Also, it is important to note that in this approach, a pair of sequences can be labeled as positive at one level and negative at another, with flexibility to select randomly.

In our special hierarchy of *Gene-Taxa* training, a notable distinction is made at the gene level, which, unlike other taxonomy levels, is not part of the natural hierarchy. Our model actively separates genes from taxonomy, enhancing its ability to differentiate between genetic characteristics and broader taxonomic classifications. Additionally, we drew inspiration from batch-hard sampling^22^ to prioritize samples and batches, detecting harder instances throughout training that exhibit the most distance between anchor-negative and anchor-positive pairs.

**Figure 8.**
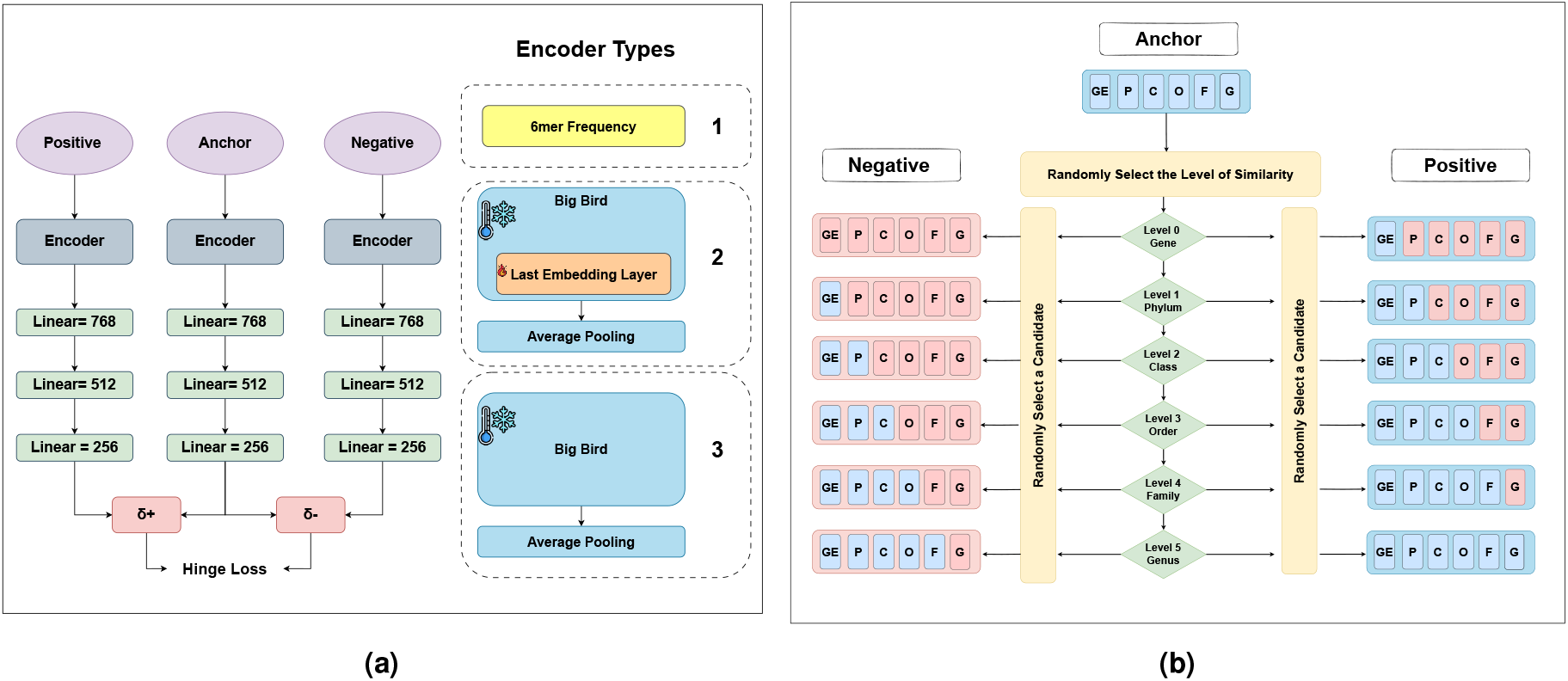
8a Model Architecture: Each branch transforms 768 (or 4096)-dimensional encoded embeddings to a 256-dimensional triplet vector. We have three types of encoders: BigBird embedding vectors, 6-mer Freq., and a model where BigBird is used with the embedding layer. 8b The diagram illustrates the hierarchical selection process for a positive and negative for an anchor in our gene-taxa dataset. First, the level of similarity is randomly determined (e.g., Ge: Gene, P: Phylum, C: Class, O: Order, F: Family, G: Genus). Positive samples match the anchor at this level, while negative samples are chosen from one level up to ensure dissimilarity.

A key improvement in our framework is its independence from the type and number of hierarchical levels. Our innovation lies in the framework’s ability to handle various hierarchies and adapt to changes in the hierarchy structure, such as adding or removing levels or incorporating new tasks. To test this, we trained the model on a promoter dataset containing just one level—whether it is a promoter or not—and it demonstrated the framework’s adaptation capabilities. We conducted extensive studies to determine the optimal batch size, number, and order of levels for hierarchical sampling. Detailed results of these studies are provided in the supplementary material.

We utilized the Margin Ranking Loss technique during our model training to optimize a novel embedding space. This approach aimed to draw anchor-positive pairs closer together, effectively reducing their distance, while simultaneously pushing anchor-negative pairs further apart, thereby increasing their Euclidean distance.

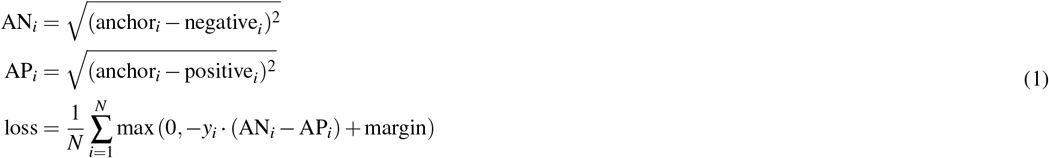

Here, AN_*i*_ represents the Euclidean distance between the anchor and negative embeddings, while AP_*i*_ denotes the Euclidean distance between the anchor and positive embeddings. The loss function loss averages the maximum of zero and the difference between the anchor-negative distance and the anchor-positive distance, weighted by the labels *y*_*i*_ and a margin parameter.

### Confidence Score

Our objective is to derive a confidence score for each prediction at every hierarchical level. To accomplish this, we employed a hybrid methodology leveraging two key techniques: I) A confidence predictor, which calculates the confidence score based on the raw distance value of the query point to the best match in the training set. II) Utilization of class probabilities derived from neighborhood embeddings specific to each query. Consider Figure 9a, which visually depicts the necessity of integrating both metrics in determining the confidence score. For instance, comparing two query points, yellow and green, which exhibit similar decision boundaries and nearest neighbors, the yellow point demonstrates a significantly lower distance to its best hit compared to the green point. Consequently, the confidence score for the yellow query should exceed that of the green query. Further, let’s analyze the scenario involving the yellow and purple stars. Although both share equidistant nearest neighbors, the surrounding point probability differs; purple records 0.4 while yellow boasts 0.6. Consequently, the confidence score for the yellow query should surpass that of the purple query. However, outliers such as the blue star, which lie considerably distant from the training points, challenge the importance of neighborhood class probability due to their significant deviation from decision boundaries. In such outliers’ cases, confidence scores may solely rely on distance to the nearest neighbor.

**Figure 9.**
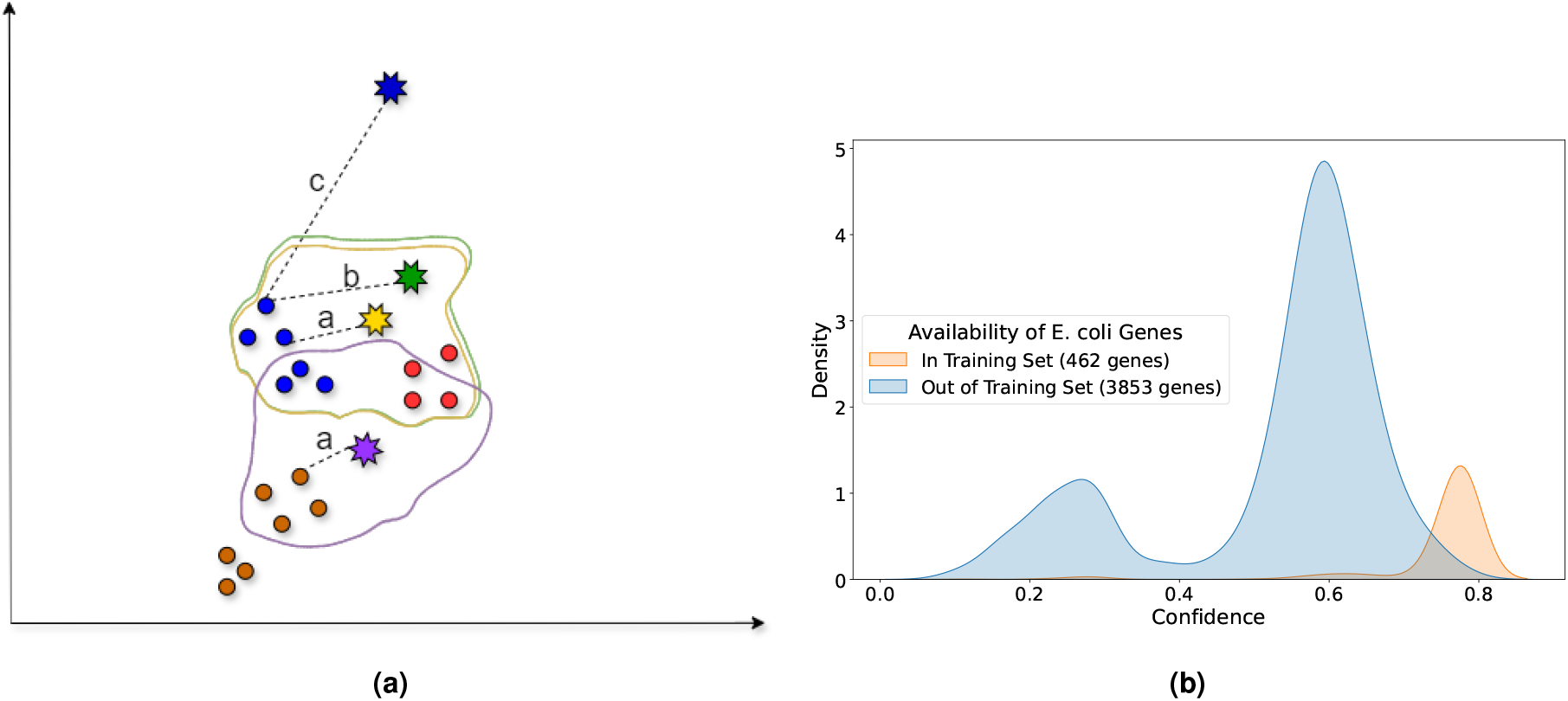
9a Confidence Score Illustration: Circular points represent the training set, with different colors corresponding to different classes, while star points represent query points. The outlines around the stars depict decision boundaries. In this example, the distance of query point *a < b* ≪ *c*, illustrating how the confidence score could vary despite the proximity of the nearest neighbors. The confidence score calculation integrates both the distance to the closest training point and neighborhood class probabilities. The blue star, despite being equidistant from its nearest neighbor as others, receives a lower confidence score due to its outlier status and the influence of decision boundaries. Figure 9b shows the KDE plot illustrating the distribution of confidence scores for genes of ***E. coli*** Genes are categorized based on their availability in the training set, with “In Training Set (**462 genes**)” indicating genes present in the training data and “Out of Training Set (**3853 genes**)” indicating genes absent from the training data. This shows the power of the confidence score as a quality estimator of predictions for users to ensure the results.

In light of these insights, we formulated a function to estimate confidence scores by harnessing the information encapsulated within neighborhood training points for each query. Let *D* represent the set of distances between query points and their closest target points in the validation set (conf_set). This conf_set includes instances from the training set but not appearing in the training set used for validation, ensuring that the distances are representative of the data distribution. Let *H* be the set of all hierarchical levels. For a specific level *ℓ* ∈ *H*, ensure that the predictions at all upper levels *u* ∈ *H* (where *u < ℓ*) are correct. To calculate the F1 score for each level, we further filter the data based on hierarchical levels. For the most upper level, such as gene, the F1 score is calculated as an accuracy F1 score. However, for lower levels, the dataset is split based on upper levels, and we should consider inner distances as reliable distances to calculate the confidence score. The dataset is further filtered to include only rows where the predictions at all upper levels are correct:

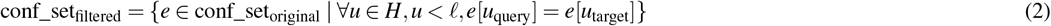

Using this filtered data, the precision and recall for the current level *ℓ* are calculated, leading to the final F1 score for the specified threshold.

For each distance threshold *t*_*i*_ in *D*, we filter the conf_set data based on the condition that the distance between query points and their closest target points is less than or equal to *t*_*i*_. This filtered dataset is used to calculate the F1 score *F*1(*t*_*i*_) for the current hierarchical level *ℓ*, considering the correctness of predictions at upper levels. The F1 score is calculated as follows:

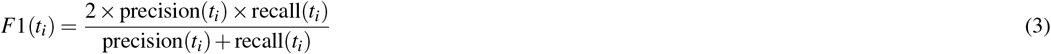

where precision and recall are defined as:

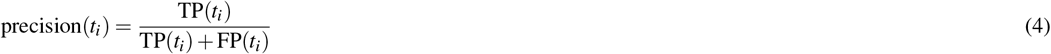

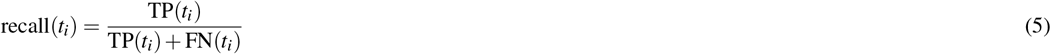

Here, TP, FP, and FN denote the true positive, false positive, and false negative counts, respectively. Once we have calculated the F1 scores for all thresholds in *D*, we use them as the target variable *y* and the corresponding distance thresholds as the input variable *X* to train a simple neural network to predict F1 based on distance threshold. This neural network, denoted as *N, N* : *t*_*i*_ → *F*1 is a function that maps distance thresholds to F1 scores: The architecture of the neural network *N* is a fully connected feedforward network with multiple hidden layers, with the output layer having a single neuron since it’s a regression model. The activation function in the hidden layers is ReLU. The neural network is trained using Mean Squared Error (MSE), which measures the difference between the predicted F1 scores and the actual F1 scores:

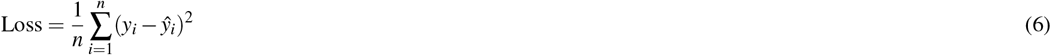

After training the model, we use it to predict the confidence score 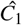 for a given distance threshold 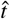, providing us with a confidence score 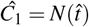. This kind of representation helps us ensure that the confidence score is bounded between 0 and 1, whereas comparing with raw distance values, which are not normalized, may not provide meaningful low and high values.

Additionally, very low F1 scores typically indicate low confidence, while high F1 scores suggest high confidence. This can be interpreted akin to a threshold finder. Next, we calculate class probabilities of target as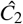, which represents the number of times class *i* (the most common class in the neighborhood) was present among the total number of nearest neighbors considered, 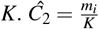, and then we compute the final confidence as follows: 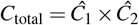.

### Dataset

#### Gene and Taxonomy Dataset

We obtained the complete Basic genome dataset using Woltka’s pipeline^27^, comprising 4634 genomes. A specific characteristic of this dataset is that only one genome is included for each genus, making it unique and challenging. After considering the taxonomic properties, we attempted to download all CDS files from the NCBI database for the Basic genome dataset. Subsequently, we extracted all coding sequences (CDS) from the Basic genomes dataset, resulting in 8 million distinct CDS. We then focused our study on bacteria and archaea, excluding genomes from viruses and fungi, which often lack sufficient gene information.

To ensure the initial annotation accuracy of genes in the dataset, we filtered out hypothetical proteins, uncharacterized protein and sequences lacking gene labels. Another issue encountered during our observation of gene labels in NCBI was the potential unreliability of gene names, possibly due to misspellings or differences in nomenclature. To address this, we retained only genes with more than 1000 samples and also imposed a filter to ensure a minimum number of sequences per phylum, considering only those with more than 350 sequences. This curation process yielded a dataset of 800,318 gene sequences, representing 497 gene types across 2,046 genera.

To assess the generalizability of our model, we deliberately constructed four types of datasets. One dataset for training, collectively referred to as the *Train Set*. Additionally, we created another dataset for testing, referred to as the *Test Set*, which comprises the same classes at all levels (same genus and same gene with *Train Set*) but with different samples. We also excluded 18 different phyla, designated as the *Taxa Out Set*, which have the same gene as the training set but from different phyla. Furthermore, we excluded 60 different genes from the *Train Set*, all originating from the same phyla, forming the *Gene Out Set*. In all testing sets, we also made sure to include only CDS that have only one representation for a genome, because we observed that once we have downloaded the CDS files for different genomes like *Hungatella hathewayi* species, we may have multiple gene sequences for one type of gene (*lepB*, for instance, has 34 representations for this species). So, we have removed such genes from our analysis because we found that in some species it may have multiple gene representations in NCBI but these genes may not be from the same species^65^. Therefore, to add more validity to our test datasets, we removed those sequences from the analysis as well. Our goal was to include holdout sets that represent diverse aspects, allowing us to evaluate the model’s performance with unseen data. The detailed information regarding the exact number of samples and the range of values per class is presented in Supplementary Table 1. Additionally, the dataset selection strategy is provided in Supplementary Figure 1.

***For the short-fragment dataset***, we extracted 400bp fragments from the 800k-sequence gene dataset. Our approach involved selecting 400bp fragments from various regions of the gene sequences, ensuring a minimum 50bp distance between them. This was done using the range ***Range:[0, Gap: 50, length(gene_sequence)]***. We believed this strategy was essential to avoid selecting fragments with minimal base-pair differences and to mimic sequences that do not necessarily start with an open reading frame.

Some of these fragments are not open reading frames (ORFs), which is significant because, in real metagenomic sequences, fragments can originate from any part of the genome and are not necessarily confined to ORFs. To address this, we utilized our curated gene dataset to ensure that the short-fragment dataset includes gene-specific information beyond just taxonomy. This approach is crucial for training the gene-taxa version of our model effectively.

#### Promoter Dataset

In this study, we utilized the promoter dataset provided by Ligeti et al.^46^ for training and testing our promoter prediction models. The promoter dataset by Ligeti et al. consists of experimentally validated promoter sequences primarily drawn from the Prokaryotic Promoter Database (PPD), which includes sequences from 75 different organisms. To ensure a balanced and comprehensive dataset, non-promoter sequences were generated using higher and zero-order Markov chains.Additionally, an independent test set focusing on *E. coli sigma70* promoters^66^ was used to benchmark the models against established datasets. The non-promoter sequences were constructed using three methods: coding sequences (CDS) extracted from the NCBI RefSeq database, random sequences generated based on a third-order Markov chain, and pure random sequences generated using a zero-order Markov chain. The balanced distribution of these non-promoter sequences (40% from CDS, 40% from Markov-derived random sequences, and 20% from pure random sequences) was crucial for thorough evaluation and robust model training. The inclusion of the independent *E. coli sigma70* test set, curated by Cassiano and Silva-Rocha (2020), further validated the effectiveness of the promoter prediction models, ensuring no overlap with the training data and providing a rigorous benchmark for model performance.

#### Antimicrobial Resistance Dataset

We utilized an integrated dataset combining the CARD^39^ v2 and MEGARes^38^ v3 datasets, referred to as the Antimicrobial Resistance Dataset, following methodologies from previous studies^40^. Classes with fewer than 15 samples were removed as they hindered the attainment of meaningful results during data splitting. The remaining data was divided into 75% for training, 20% for testing, and 5% for validation. After integrating the data using the EBI (European Bioinformatics Institute) ARO (Antibiotic Resistance Ontology) ontology search, it was similarly divided. Classes that yielded non-meaningful results were also excluded. The MEGARes dataset comprises 9733 reference sequences, 1088 SNPs, 4 antibiotic types, 59 resistance classes, and 233 mechanisms. The CARD dataset includes 5194 reference sequences, 2005 SNPs, 142 drug classes, 331 gene families, and 10 resistance mechanisms. The EBI ARO ontology provides hierarchical group information for genes, allowing gene family class information to be integrated into a higher-level hierarchy. For MEGARes, there are 589 gene family text information classes, while CARD has 331. There are 300 and 166 datasets with only one sample in their respective gene family classes for MEGARes and CARD, respectively.Resistance mechanisms categories are integrated based on the 6 categories of CARD. The original 8 categories were reduced to 6 by excluding various class combinations and those with very few samples. Drug classes are integrated using 9 common drug classes found in competing models. The integration is based on names and theories and has been validated by experts in the field.

### FAISS

In our framework, FAISS (Facebook AI Similarity Search)^26^ played a pivotal role as a cornerstone element for conducting similarity search tasks. This versatile library, designed by Johnson et al. (2019)^26^, specializes in facilitating approximate nearest neighbor search (ANNS) on vector embeddings, addressing various domains and applications. Leveraging FAISS, we efficiently conducted similarity searches on our collection of query embeddings, benefiting from its indexing techniques that involve preprocessing, compression, and non-exhaustive search methods detailed in Supplementary Table 4. These strategies enabled swift retrieval of nearest neighbors, whether based on Euclidean distance or highest dot product^67^. This streamlined approach greatly aided in identifying similar embeddings within our expansive dataset while effectively managing computational resources and memory overhead.Additionally, FAISS provides optimized versions for both CPU and GPU platforms^64^ with the latter proving particularly advantageous, especially when dealing with high-dimensional vectors exceeding 1000 dimensions. This GPU acceleration, noted for its significant performance boost, accelerated our similarity search tasks, especially vital for processing large-scale datasets, leveraging the parallel processing capabilities inherent to GPUs.

### Pre-training the BigBird Model

In this study, we utilized the BigBird model, a transformer-based architecture specifically designed to handle long sentences, to represent our gene sequences. The BigBird model enhances the standard transformer model by incorporating sparse attention mechanisms, allowing it to efficiently manage much longer contexts, which is particularly advantageous for genomic data characterized by lengthy sequences and complex dependencies^25^. We follow the approach in MetaBERTa^17^ to select the parameters for the BigBird model. We trained the BigBird model using a sequence length of 4096, with a batch size of 16, and an embedding dimension of 768. The feed-forward neural network within the transformer layers was configured with a dimension of 3072. The model employed 4 attention heads and comprised 4 transformer layers, facilitating the learning of hierarchical representations. The Adam optimizer with an epsilon value of 1e-8 was used for training. Training was conducted over 2 epochs. The vocabulary size of our model is based on 6-mer, which corresponds to 4096 tokens. This selection can be attributed to the fact that each 6-mer represents two codons, which correspond to amino acids, ensuring more functional information^17^. These configurations were selected to optimize the model’s performance for genomic sequence analysis, leveraging its unique capabilities to manage and learn from long sequences efficiently. Our BigBird model, MetaBERTa-bigbird-gene, is available at https://huggingface.co/MsAlEhR/MetaBERTa-bigbird-gene.

### Benchmarking and Configuration of Comparative Tools

All benchmark tools were trained and evaluated on a standardized dataset to maintain consistency in comparisons. The configurations applied to each tool are outlined below.

**Kraken 2** (gene-taxa classification): Kraken 2 was configured with `-minimum-hit-groups 1` and `-confidence 0`, enabling a more comprehensive search to increase sensitivity for unclassified taxa.

**MMseqs2** (AMR and gene-taxa classification): The `mmseqs easy-search` command was executed with parameters `-max-accept 1, -max-seqs 1`, and `-search-type 3` to focus on the best-hit nucleotide-to-nucleotide alignment.

**BLASTN** (AMR identification): The `blastn` tool was used with the following arguments: `-db ./myblastdb/triplet, -evalue 0.01, -word_size 11`.

**BERTax** (gene-taxa classification): The pre-trained BERTax model was used without further fine-tuning. Access to the tool is available at https://github.com/rnajena/bertax.

**ABRicate** (AMR identification):ABRicate was indexed with a custom AMR database to perform antimicrobial resistance (AMR) gene detection. Further details are available at https://github.com/tseemann/abricate.

**DeepMicrobes** (gene-taxa classification): We trained DeepMicrobes to adapt for long-read data, with separate training at both the family and gene levels. The model was configured for gene-level classification specifically for this purpose, in addition to family-level classification. The full methodology follows their original approach, with details available at https://github.com/MicrobeLab/DeepMicrobes, using the following arguments: `-model_name attention, -vocab_size 32898, -train_epochs 10, -batch_size 256, -max_len 2048`.

## Acknowledgements

This work is supported by National Science Foundation under Grant Numbers #1936791, #1919691 and #2107108. We thank the University Research Computing Facility for their paid services.

## Data availability

All data used in this manuscript originates from publicly available databases. The sequence data used for pre-training and Scorpio of Gene-Taxa are available in the Zenodo repository https://zenodo.org/records/12964684. Additionally, the dataset for promoter detection was downloaded from the ProkeBERT repository on Hugging Face, available https://huggingface.co/neuralbioinfo.

## Code availability

The source code for the paper is freely available online under an open source license https://github.com/EESI/Scorpio.

## Author Contributions

M.S.R. conceived and designed the study, developed the algorithms and software, analyzed the data, interpreted results, wrote the manuscript, and prepared figures. B.A.S. conceived and designed the study, interpreted the data, analyzed the data, and reviewed the manuscript. J.C.M. designed the study and interpreted the data. J.R.B. designed the study, analyzed the data, interpreted results, and reviewed the manuscript. H.Y. prepared the data and reviewed the manuscript, G.H. ran the software and reviewed the manuscript. G.L.R. conceived and designed the study, interpreted results, analyzed the data,supervised the work, and wrote the manuscript.

