## Supplemental Information for "Scorpio : Enhancing Embeddings to Improve Downstream Analysis of DNA sequences"

##### *Dataset Selection Strategy*

The selection of training and test sets, including the Test set, Taxa Out set, and Gene Out set, follows a carefully designed strategy that ensures balanced representation and hierarchical relevance within a dataset containing one genome per genus. This approach considers both gene presence within genera and taxonomic distinctions.Referring to the Figure 1, the dataset partitioning strategy is as follows:

**Test Set:** When a gene (e.g., Gene 1 from Genome 1) is included in the test set, another gene from the same genome (such as Gene 2) is placed in the training set. Additionally, similar genes from other genomes within the same genus are included in the training set. This structure ensures that the model is exposed to the genus and gene type, even if it hasn't seen the exact test sequence, allowing for genus-level generalization.

**Taxa Out Set :** For the Taxa Out set, genomes from different phyla, such as Genome 4 from Phylum 2 (illustrated in the figure), are entirely excluded from the training set. However, the training set still contains genomes from other phyla (e.g., Phylum 1), providing indirect exposure to related gene types without direct access to the excluded taxonomy. This ensures that while the specific phylum is omitted, the model learns from similar genes in related taxonomic groups.

**Gene Out Set**: In the Gene Out set, specific genes (e.g., Gene 3) are entirely excluded from the training set, while other genes (e.g., Gene 1 and Gene 2) from the same genomes are retained. This approach enables the model to gain insights into the genomic context without direct exposure to the excluded gene, supporting gene-level generalization.

This dataset selection strategy optimizes the model's ability to generalize across hierarchical taxonomic levels and gene variations, enhancing its robustness in diverse scenarios.

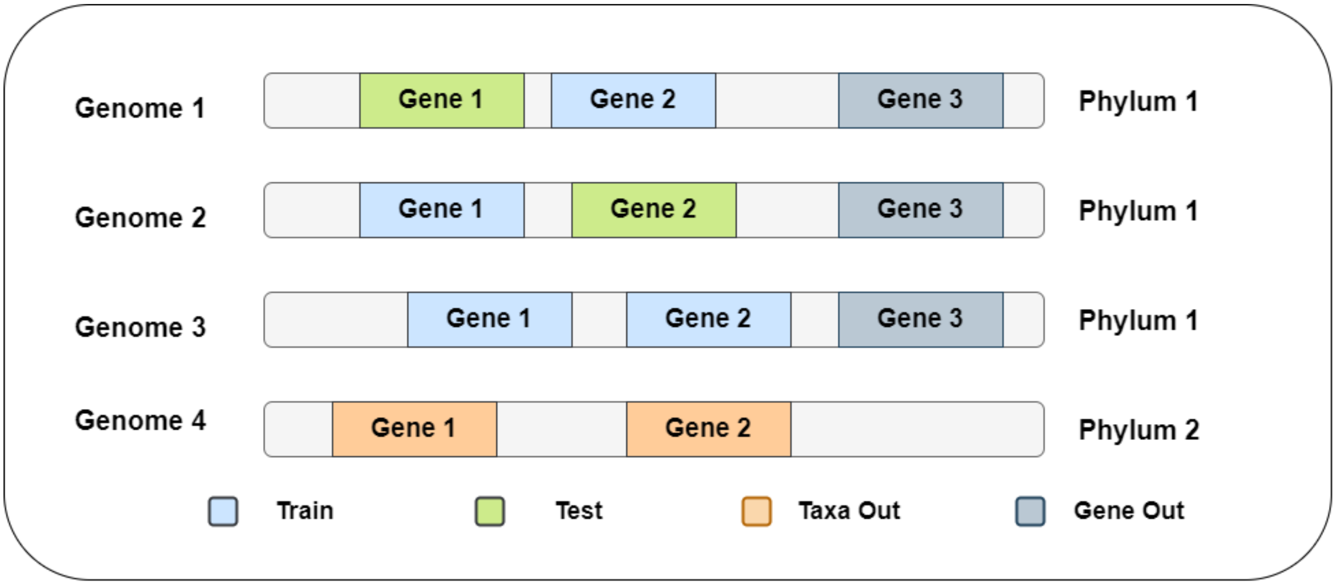


**Figure 1.** Illustration of genome and gene selection strategy for training and test sets. For the test set, a genome (e.g., Genome 1) may have a gene included in the training set, ensuring that the model has prior knowledge of genes from the same genome. For the Taxa Out set, genomes from different phyla (e.g., Genome 4) are excluded, but their genes (Gene1 , Gene2) are still represented in the training set. For the Gene Out set, specific genes (e.g., Gene 3) are excluded while the rest of the genome's genes remain in the training set.

**Table 1.** Distribution of the Gene-Taxa Dataset (Full Gene Dataset) Across Various Taxonomic Levels. This table provides a detailed breakdown of the number of classes and the range of # of iexamples in each class for each taxonomic level in the training set, test set, taxa-out set, and gene-out set. The dataset spans multiple hierarchical levels, including gene, kingdom, phylum, class, order, family, and genus. The ranges of # of examples in each class indicate the minimum and maximum instances found in a classwithin each taxonomic level for the respective sets. The total number of instances for the training set, test set, taxa-out set, and gene-out set are also provided.

| Level | Train set | | Test set | | Taxa out | | Gene out | |
|---|---|---|---|---|---|---|---|---|
| | #Classes | Instance # Range | #Classes | Instance # Range | #Classes | Instance # Range | #Classes | Instance # Range |
| gene | 437 | 2378 > x > 787 | 437 | 500 > x > 207 | 436 | 39 > x > 1 | 60 | 985 > x > 140 |
| kingdom | 2 | 533587 > x > 13936 | 2 | 137061 > x > 3463 | 1 | x = 11800 | 2 | 42089 > x > 1328 |
| phylum | 27 | 228975 > x > 578 | 27 | 59203 > x > 154 | 18 | 1778 > x > 298 | 27 | 21455 > x > 59 |
| class | 83 | 92784 > x > 64 | 83 | 24108 > x > 13 | 25 | 1289 > x > 274 | 83 | 9055 > x > 7 |
| order | 204 | 32643 > x > 53 | 204 | 8219 > x > 13 | 27 | 1079 > x > 165 | 204 | 2681 > x > 5 |
| family | 505 | 18713 > x > 53 | 505 | 4776 > x > 13 | 33 | 723 > x > 149 | 505 | 1582 > x > 4 |
| genus | 1929 | 391 > x > 49 | 1929 | 111 > x > 6 | 39 | 370 > x > 103 | 1928 | 45 > x > 1 |
| Total | | 547523 | | 140524 | | 11800 | | 43417 |

##### *Effect of Order and number of Hierarchical levels*

Hierarchical Triplet Training is highly dependent on the order and number of hierarchical levels used. This dependence arises from the triplet set (Anchor, Positive, Negative) selection process, which is based on hierarchical structure and similarity levels for positive and negative samples. The hierarchical structure significantly impacts the accuracy and F1-macro scores of different levels. Supplementary Table 2 showcases performance metrics across different hierarchical arrangements, revealing that the number of levels and their order can substantially affect classification outcomes. In all these models, we keep all the model parameters the same (including the model architecture, batch size, and number of epochs) and only change the hierarchy's structure and number of levels to examine their performance. We observe that starting with the Phylum level (phylum-first model) as the highest hierarchy leads to higher accuracy and F1 scores compared to other training types in taxonomic levels. Conversely, choosing Gene as the first level results in higher performance in gene classification for that model compared to starting with the phylum-first model. This phenomenon aligns with the embedding perspective and representation learning of hierarchical levels, where embeddings are first clustered based on the highest level of hierarchy and then further refined into inner clusters based on how the hierarchy is arranged. Figure 7 illustrates this phenomenon, showing that the t-SNE results from the gene-first model indicate that embeddings first cluster based on genes and then form additional inner clusters within each gene cluster. This results in higher accuracy at the gene level due to the higher number of nearest neighbors for genes compared to other taxonomic levels. Another interesting point we tested is the effect of the number of hierarchical levels on model performance. Here, we compared models trained on six levels versus two levels of hierarchy. We observe that having more levels of hierarchy helps improve performance. This improvement likely stems from the model's ability to better understand the overall hierarchical features, aiding in the selection of positive and negative samples based on varying similarity levels, which better matches phylogenetic information and clusters them more effectively.

**Table 2.** Performance Metrics for Triplet Model for Different Hierarchical Arrangements

| | Accuracy (%) | | | | | F1-macro (%) | | | | |
|---|---|---|---|---|---|---|---|---|---|---|
| Hierarchy of Levels | Phylum | Class | Order | Family | Gene | Phylum | Class | Order | Family | Gene |
| Phylum → Class → Order → Family → Genus → Gene | 80.8 | 70.6 | 55.7 | 45.7 | 54.3 | 62.8 | 51.3 | 45.7 | 39.6 | 53.2 |
| Gene → Phylum → Class → Order → Family → Genus | 75.6 | 64.5 | 47.7 | 36.9 | 75.0 | 52.5 | 40.4 | 35.1 | 29.7 | 75.9 |
| Gene → Genus | 69.2 | 58.5 | 44.4 | 35.6 | 75.6 | 47.5 | 37.3 | 33.5 | 29.1 | 77.0 |
| Genus → Gene | 75.4 | 65.0 | 49.6 | 40.4 | 39.5 | 55.6 | 44.3 | 39.3 | 34.5 | 37.8 |

#### Evaluation on ART-Simulated Illumina 150bp Reads

In this study, we utilized 400bp fragments to enhance the contextual information around genes, supporting functional and taxonomic analyses in metagenomic studies. Certain next-generation sequencing (NGS) platforms, such as the Ion Torrent Genexus (Thermo Fisher) and the Illumina MiSeq, are capable of producing read lengths of up to 400bp in targeted sequencing workflows. To provide a broader comparison with commonly used short-read lengths, we also simulated 150bp reads using the Illumina ART simulator. The parameters used included `-ss HS25` (HiSeq 2500), `-f 20` (20× coverage depth), and

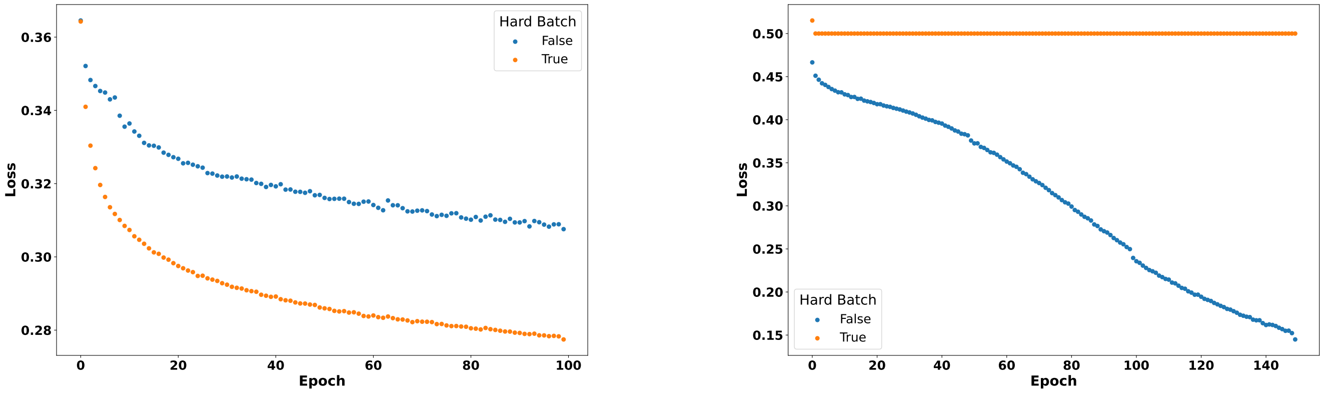


**(a)** Gene-taxa dataset: Batch-hard sampling

**(b)** Promoter dataset: Batch-hard sampling

**Figure 2.** The plots compare the effect of the hard-batch effect on triplet training for the gene-taxa dataset and the promoter dataset. We observe that the impact of using the batch-hard sampling effect varies depending on the dataset. In the promoter dataset, it seems that using the hard-batch effect reduces the learning ability of the model. In contrast, in the gene-taxa dataset, the batch-hard sampling effect helps improve training. This difference could be due to the nature of the hierarchy and the number of levels, which is greater in the gene-taxa dataset compared to the promoter dataset.

error-related configurations: `-ir 0.00009`, `-dr 0.00011` (insertion and deletion rates for the first read), and `-ir2 0.00015`, `-dr2 0.00023` (for the second read). These settings closely replicate typical Illumina sequencing conditions.

Table 3 presents the performance of our models, Scorpio-BigDynamic, Scorpio-BigEmbed, and BigBird, in comparison to Kraken2. The evaluations were conducted on ART-simulated datasets comprising 2.9 million reads, with the training dataset consisting of gene sequences of an average length of 4096bp.

**Table 3.** Comparison of Accuracy and F1-macro scores across taxonomic levels for Scorpio-BigDynamic, Scorpio-BigEmbed, and Kraken2. Each model was trained on the gene-taxa dataset and tested on simulated reads generated using ART with realistic sequencing parameters.

| | Accuracy (%) | | | | | | F1-macro (%) | | | | | |
|---|---|---|---|---|---|---|---|---|---|---|---|---|
| **Level** | **Kingdom** | **Phylum** | **Class** | **Order** | **Family** | **Genus/Species** | **Kingdom** | **Phylum** | **Class** | **Order** | **Family** | **Genus/Species** |
| **Scorpio-BigDynamic** | 96.0 | 25.5 | 12.1 | 3.0 | 1.0 | 0.1 | 49.9 | 4.6 | 1.5 | 0.6 | 0.2 | 0.0 |
| **Scorpio-BigEmbed** | **97.2** | 27.9 | 14.8 | 3.0 | 1.1 | 0.1 | 50.4 | 4.8 | 1.6 | 0.6 | 0.2 | 0.1 |
| **BigBird** | 95.7 | **41.3** | **23.0** | **8.7** | **4.3** | **1.4** | **59.6** | **11.9** | 6.0 | 3.9 | 2.5 | **1.3** |
| **Kraken2** | 0.36 | 0.15 | 0.08 | 0.02 | 0.01 | 0.005 | 56.3 | 11.4 | **6.7** | **4.0** | **2.7** | 1.2 |

Our results suggest that Scorpio generally achieves higher accuracy across all taxonomic levels compared to Kraken2, which classified only 11,000 reads, representing about 0.38% of the sequences. However, Scorpio's lower F1-macro scores may reflect sensitivity to sequencing errors, particularly in underrepresented classes. This indicates that Scorpio might face challenges with sequencing errors, contextual inaccuracies, or the disparity between the 150bp reads used for testing and the average 4096-length gene embeddings used during training. These factors likely contribute to occasional misclassifications. While Scorpio demonstrates strong performance in dominant classes, its lower precision in underrepresented classes impacts the overall F1-macro score, highlighting areas for improvement in handling diverse and error-prone reads. BigBird, on the other hand, consistently achieves higher accuracy across almost all taxonomic levels, except at the kingdom level. Its performance is particularly robust at intermediate levels, such as phylum and class, and it also achieves higher F1-macro scores at the phylum, kingdom, species, and genus levels. These results emphasize BigBird's ability to generalize effectively across diverse taxonomic levels, making it better suited for tasks involving complex sequencing data and varying read lengths.

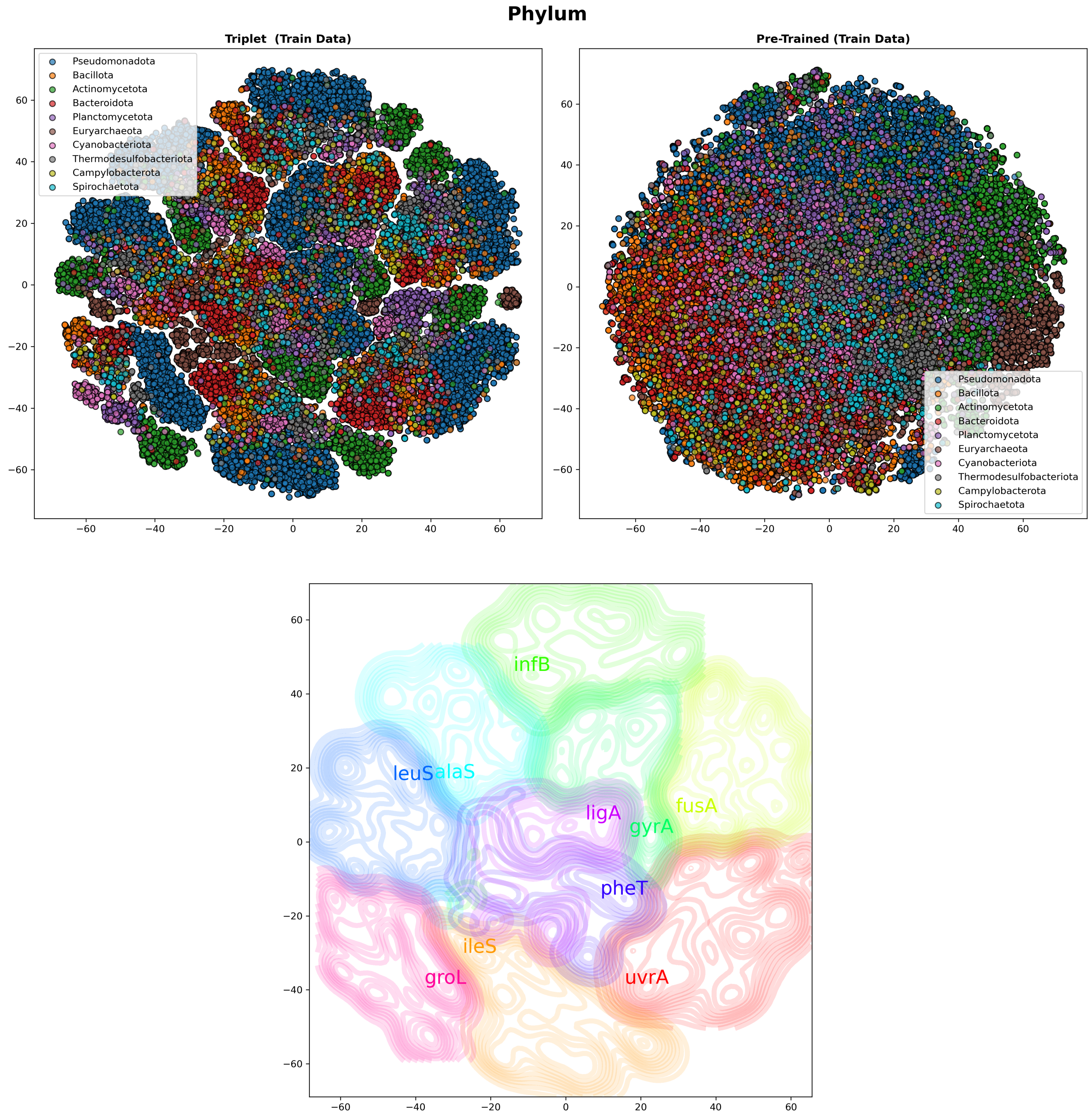


**Figure 3.** (a) t-SNE visualization of phylum embeddings colorized based on the 10 most common phyla in the dataset using the Scorpio-BigEmbed(Triplet) and BigBird(Pre-Trained) Models. The left plot shows embeddings from the Scorpio-BigEmbed model, while the right plot shows embeddings from the BigBird model. We observe that the Scorpio-BigEmbed model produces more distinct clusters for different phyla compared to the BigBird model. This distinction highlights the effectiveness of the Scorpio-BigEmbed model in capturing taxonomic relationships. (b) Kernel density estimation of gene embeddings for Scorpio-BigEmbed. The plot shows Gaussian distributions for each gene, with inner Gaussian distributions representing different taxa types. This visualization helps in understanding how different genes cluster together and how these clusters correspond to various taxa. By comparing this plot with the t-SNE plot, we can observe the underlying distribution patterns of gene embeddings and their taxonomic affiliations.

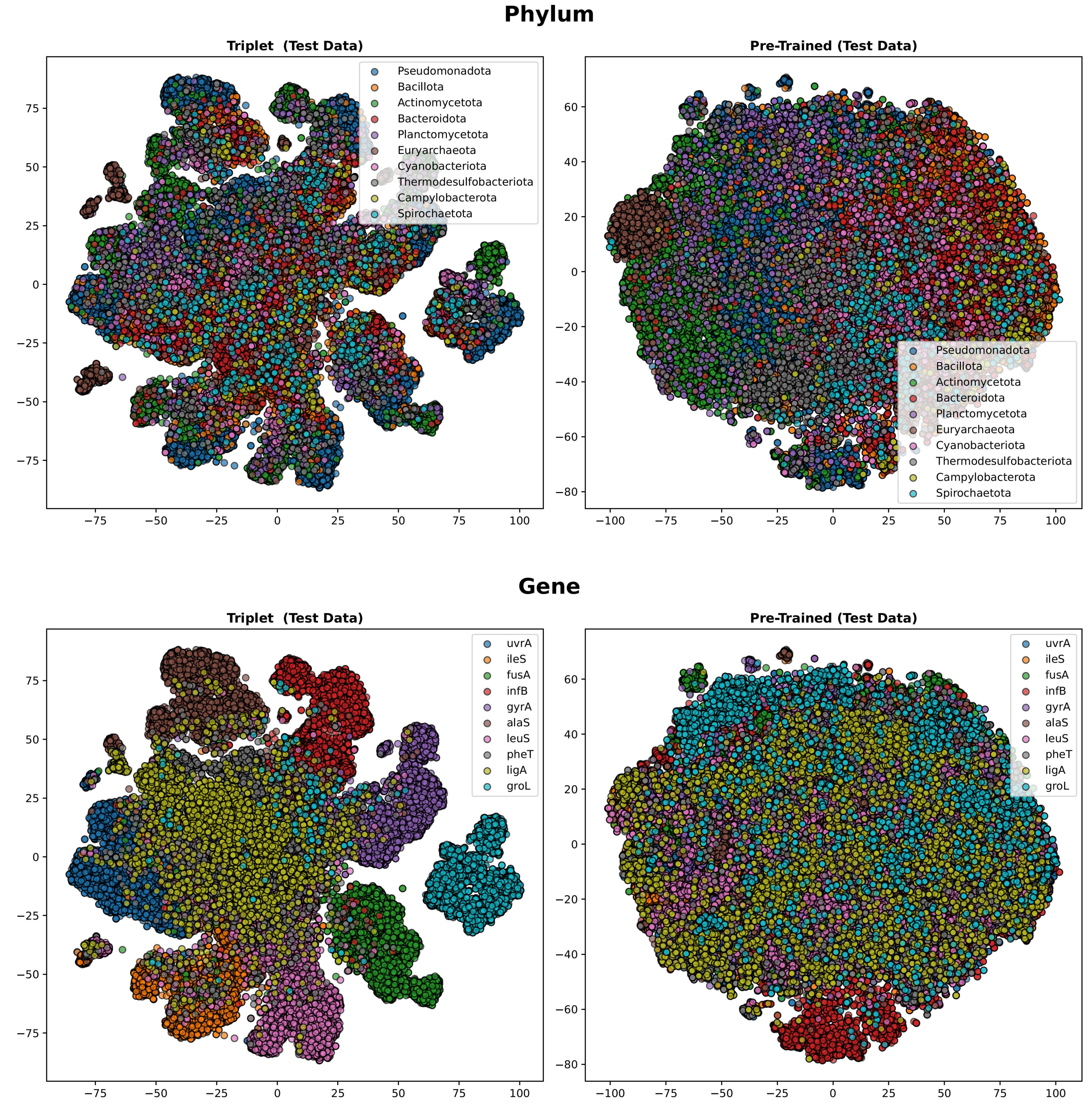


**Figure 4.** t-SNE visualization of embeddings generated by the Scorpio-BigEmbed (Triplet) and BigBird (Pre-Trained) models for test set. Plot (a) shows embeddings compared based on phylum, colored by the 10 most common phyla in the dataset. Plot (b) shows embeddings compared based on gene type, colored by the 10 most common genes. These visualizations, produced on the test set, demonstrate the generalization ability of the models. The Scorpio-BigEmbed model shows more distinct clusters for each gene, highlighting the improved clustering ability of embeddings derived from this model as compared to the pre-trained BigBird model. This clustering is also evident within each gene for the phyla clusters.

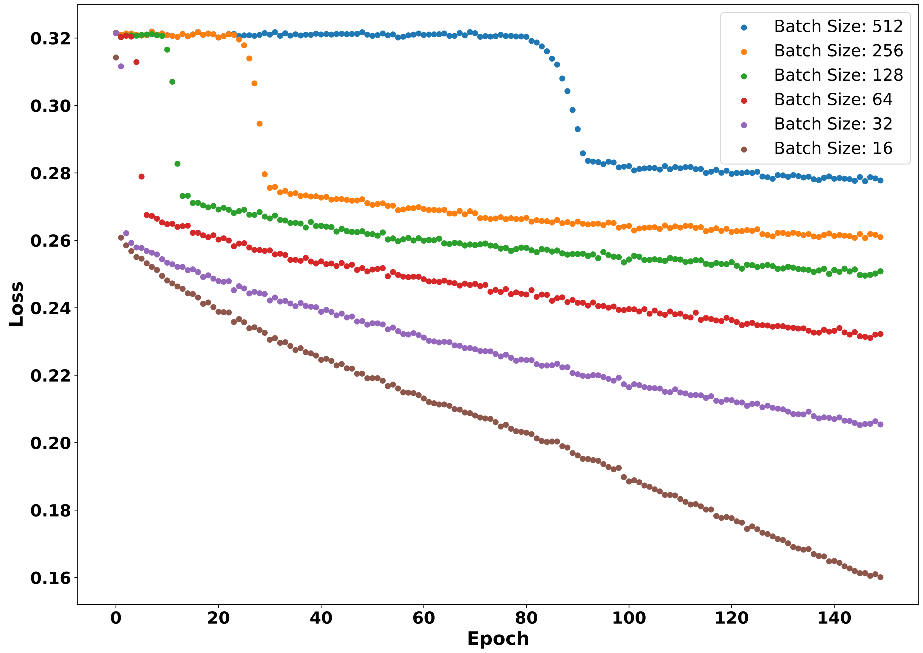


**Figure 5.** Loss vs Epoch for Different Batch Sizes: This plot illustrates the training loss against the number of epochs for various batch sizes using the Scorpio-BigEmbed model. The model leverages triplet loss in conjunction with a pre-trained Bigbird embedding to enhance the representation learning for short read gene sequences. All the experiments follow the same architecture, with the only change being the batch size, and the hard_batch parameter was set to true.

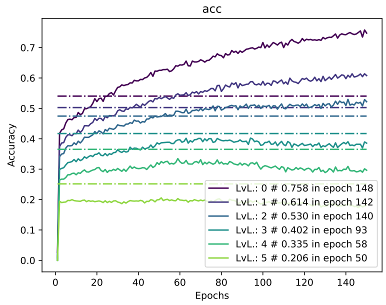


**(a)** Accuracy with Batch Size 16

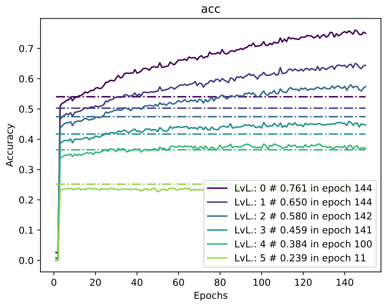


**(b)** Accuracy with Batch Size 32

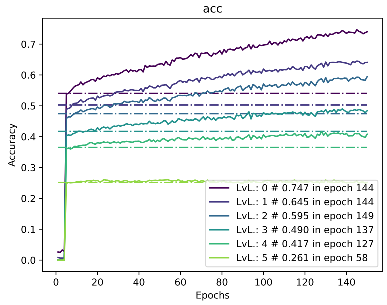


**(c)** Accuracy with Batch Size 64

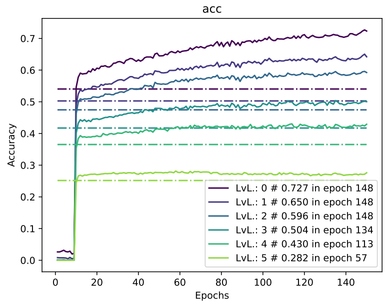


**(d)** Accuracy with Batch Size 128

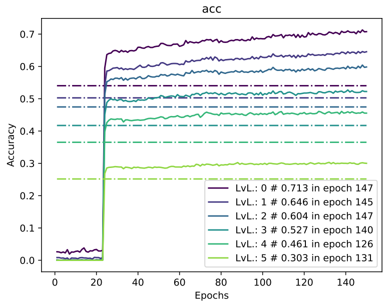


**(e)** Accuracy with Batch Size 256

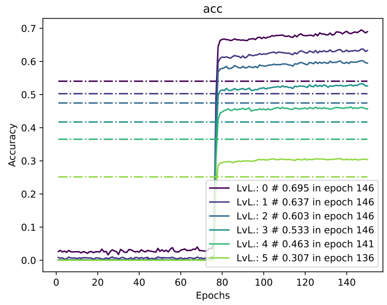


**(f)** Accuracy with Batch Size 512

**Figure 6.** Effect of Batch Size on Accuracy: This is the accuracy of Scorpio-BigEmbed on validation set during training vs epoch for different hierarchical levels (0: gene, 1: phylum, 2: class, 3: order, 4: family, 5: genus). As observed, all batch sizes surpass the baseline accuracy, but lower batch sizes converge sooner. All results are with the hard batch effect.

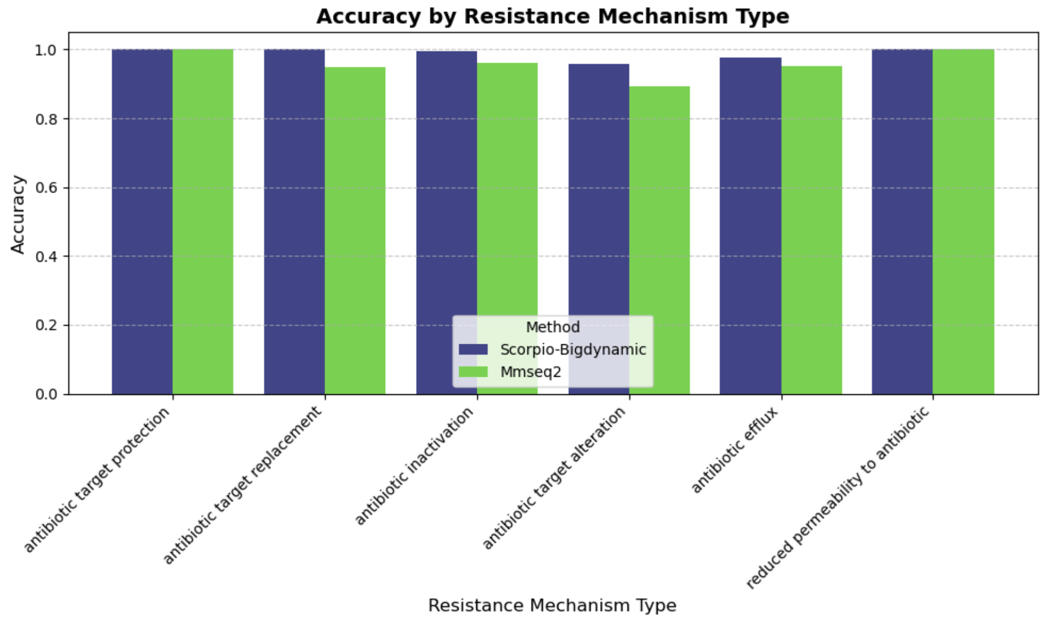


**Figure 7.** Comparison of the accuracy of MMseq2 and ScorpioBigDynamic for result mechanism types. Notably, ScorpioBigDynamic consistently performs well. The model's advantage lies more in antibiotic target alteration, where it predicts with 7% higher accuracy than MMseq2.

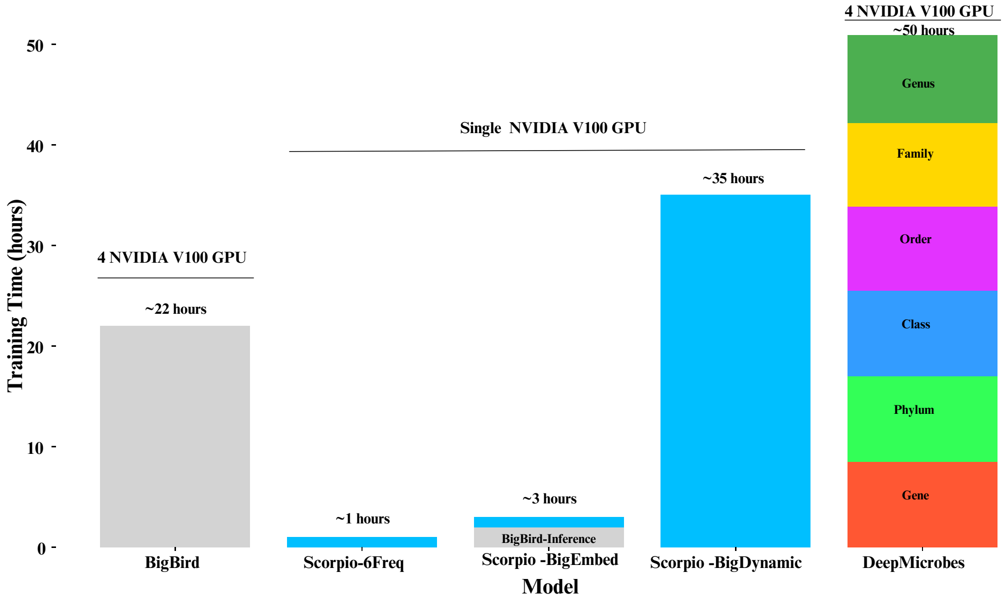


**Figure 8.** Training Time Comparison of Different Models on Various Hierarchical Levels. This demonstrates Scorpio Model's Flexibility: Unlike traditional supervised models, Scorpio employs a multi-output model that can train and gather information in a single training loop, showcasing superior generalization and multitasking capabilities. Our framework is faster, leveraging the benefits of contrastive learning with a minimal training architecture.

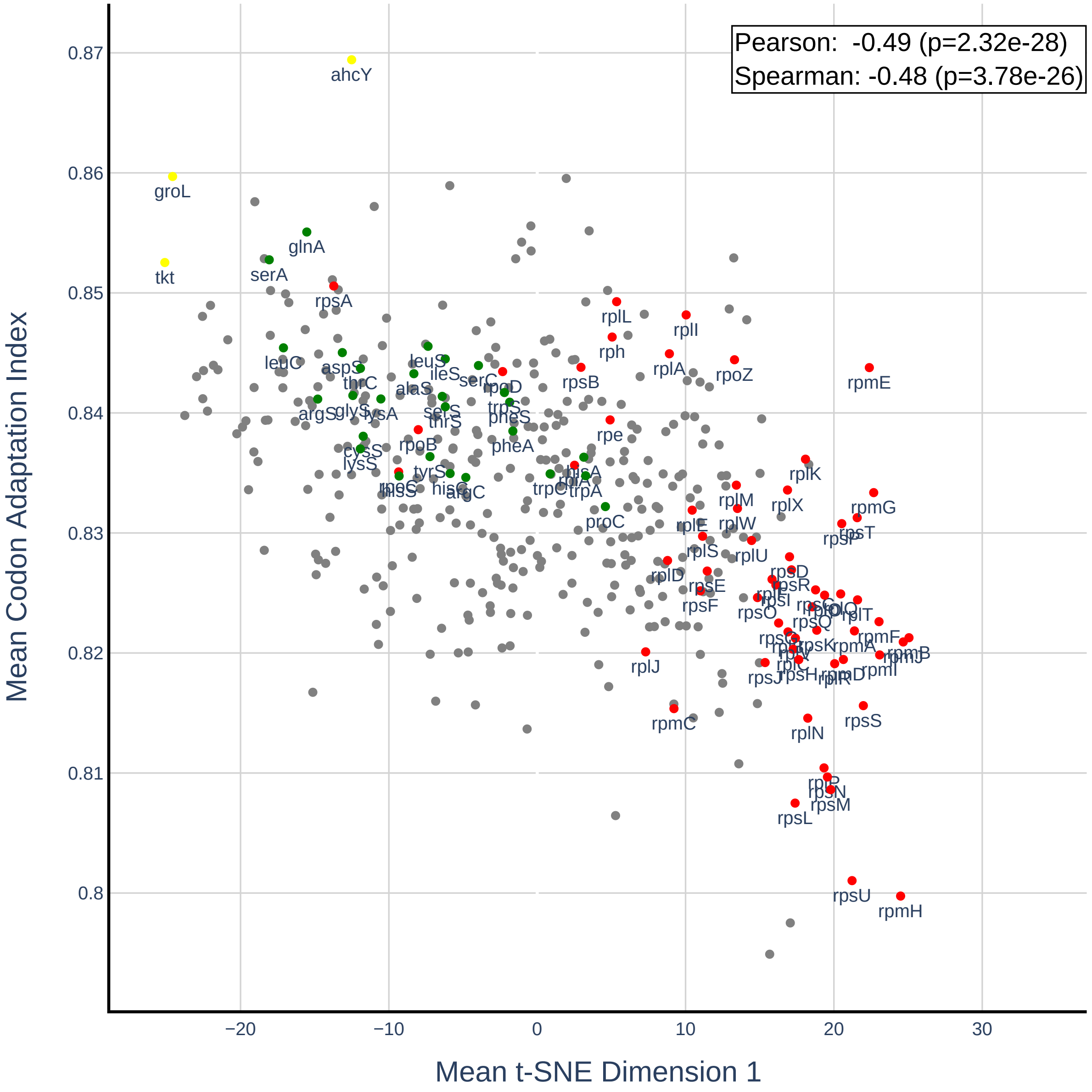


**Figure 9.** This Figure illustrates the correlation between the first dimension of the t-SNE representation and the Codon Adaptation Index (CAI) for the entire gene dataset. The plot includes all genes, with two common categories in our dataset: ribosomal proteins and aminoacyl-tRNA synthetases. The Pearson correlation between the t-SNE dimension 1 and CAI is -0.49 (p=2.32e-28), and the Spearman correlation is -0.48 (p=3.78e-26), both indicating a moderate negative correlation. Red-colored genes represent ribosomal proteins (rp*), while green-colored genes represent aminoacyl-tRNA synthetases. It is evident from the plot that these genes tend to cluster together, with ribosomal proteins forming distinct clusters and aminoacyl-tRNA synthetases forming another.

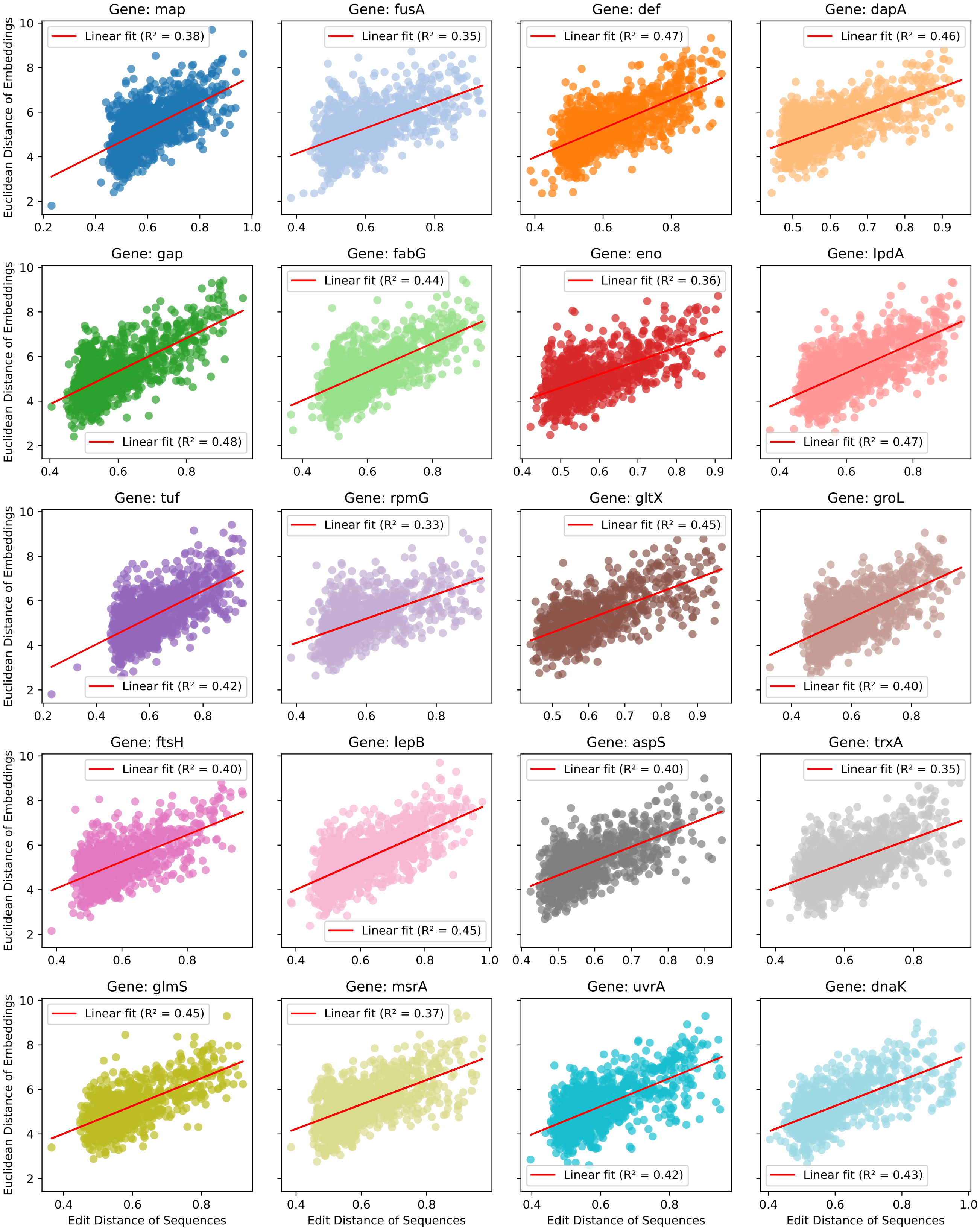


**Figure 10.** Scatter plots showing the relationship between the Euclidean distance of gene embeddings and the Edit Distance of sequences for various genes. Each plot includes a linear fit line based on a single predictor (Euclidean distance), with the corresponding $R^2$ (coefficient of determination) value indicating the strength of the linear relationship.

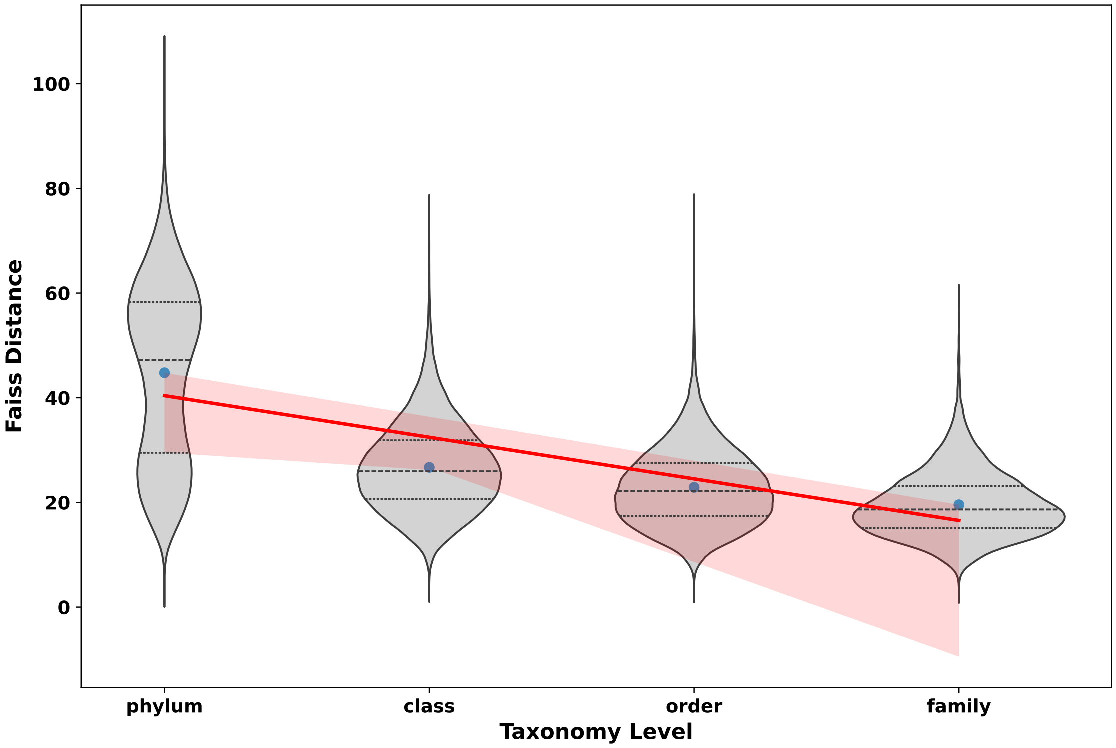


**Figure 11.** Within-taxon distribution of pairwise embedding distance for the `uvrA` gene. The violin plots depict the distribution of Faiss distances across different taxonomy levels: phylum, class, order, and family. Each plot includes quartile markers and median values. The red linear regression line, accompanied by a shaded confidence interval, highlights the trend of mean distances across the taxonomy levels. An R-squared value of 0.83 indicates a strong linear relationship between the taxonomy levels and the mean Faiss distances within-taxon. This analysis is crucial to pick the right threshold and explains why our framework introduces multiple confidence scores for prediction, allowing each hierarchical level to have its own distance threshold

### Comparative Analysis of Searching Methods

#### 0.1 ScaNN

ScaNN (Scalable Nearest Neighbors)[1] is a high-performance similarity search algorithm developed by Google, designed for efficient handling of large-scale vector searches. It leverages advanced techniques such as product quantization (PQ), anisotropic loss functions, and vector quantization-based trees to enhance both speed and accuracy in data retrieval tasks. Unlike traditional brute-force methods, ScaNN employs an asymmetric distance calculation, which quantizes only the database vectors, resulting in better distance estimates and significantly faster search times. This optimization, combined with Single Instruction Multiple Data (SIMD) in-register lookup tables, makes ScaNN highly efficient in terms of CPU and memory usage.

#### 0.2 Experiment Overview

In the comparative analysis of FAISS and ScaNN, the experiment focused on evaluating the performance of these two similarity search algorithms using a dataset of genetic sequence embeddings. These embeddings were created using a language model, enabling the comparison of similarity searches between FAISS and ScaNN. The experiment aimed to determine which algorithm performs better in terms of accuracy, sensitivity, specificity, indexing time, and search time efficiency. FAISS and ScaNN were both used to generate distance and index matrices from the genetic sequence embeddings. Various indexing methods and parameters were tested for FAISS, including Flat, IVF, and product quantization (PQ), while ScaNN was tested with pure brute force scoring, asymmetric hashing (AH), and partitioning combined with AH. The BigBird model was utilized to generate the embeddings, providing a robust basis for similarity searches and novelty detection. After parameter tuning both methods, we selected the Flat index for FAISS and for ScaNN, the optimal pipeline was found to be as follows: Build the index with 10 iterations and a dot product distance metric. Then partition the database with 2000 leaves Then utilize asymmetric hashing, with the recommended dimensions per block = 2 and anisotropic quantization threshold = 0.2. The experiment also included the classification of sequences to identify combinations of test sets, which included both test and Taxa_out gene sequences.The comparison (Figure 12) illustrates the performance disparities between the FAISS and ScaNN methods across various metrics. In terms of accuracy, FAISS exhibits superiority, boasting a higher percentage (35.4%) compared to ScaNN (31%). This trend extends to specificity and sensitivity, where FAISS outperforms ScaNN with percentages of 93.7% and 92.9% for specificity, and 92.9% and 92.5% for sensitivity, respectively. This is the largest advantage that FAISS has over ScaNN.

Examining resource utilization during indexing, FAISS generally utilizes more resources than ScaNN, which displays lower CPU usage (3.6%) compared to FAISS (10.9%). FAISS and ScaNN also showcase similar efficiency in memory usage during indexing, utilizing 678.3MB in contrast to ScaNN's 646.92MB. The story is significantly different, however, when it comes to runtime. When creating the index, FAISS excels in timing, completing the process in 1.6 seconds, whereas ScaNN takes significantly longer at 240.9 seconds. However, this trend is reversed during the searching phase, where ScaNN demonstrates its advantage with lower CPU usage (20%) compared to FAISS (62.8%). ScaNN also demonstrates efficiency in memory usage (55.798MB) and timing (2.1 seconds) during searching, even outperforming FAISS, which requires 58MB and 8.177 seconds, respectively.

It seems like ScaNN could have better results in terms of speed, memory, and CPU usage during search time. However, because we want to have parameter selection in this version of Scorpio based on performance on downstream tasks and also the possibility of searching with FAISS on GPUs, which makes it much faster than CPU search, we continue with FAISS. However, our pipeline has the option to run on both, and in future versions, we may reconsider this decision.

#### 0.3 Parameter Tuning Result of FAISS

In our evaluation of FAISS[2], we tested various indexes and parameters to assess their speed and memory usage. We employed the indexfactory() function to test multiple indexing methods. As summarized in Table4, the indexes evaluated included the base Flat index, IVF indexes, and product quantization. The Flat index, the simplest computational method provided by FAISS, employs brute force techniques to calculate distances and store vectors in a flat array. While this method typically delivers exact results, it demands significant memory and CPU resources. We also evaluated IVF indexes, which cluster database vectors using a user-defined coarse quantizer, thus reducing the number of vectors compared to the query. This approach is more efficient for larger datasets as it minimizes memory usage. Additionally, we tested product quantization (PQ), which divides the input vector into sub-vectors and processes them independently. This method significantly reduces memory usage at the expense of some accuracy. Notably, ScaNN is essentially an optimized version of the PQ indexing method in FAISS, as highlighted by authors[2]. To further understand the impact on accuracy and performance, we incorporated various preprocessing options and quantizers with IVF and PQ indexing methods. These included PCA, which reduces the dimensionality of the database matrix to enhance processing speed, and OPQ, which rotates input vectors to improve PQ indexing accuracy. Throughout this evaluation, we maintained consistent hyperparameters to provide an unbiased comparison of each indexing method's performance metrics.

The results, presented in Table 4, indicate that the Flat index excels in accuracy and speed, as expected due to its simplicity. However, its high memory and CPU requirements make it less suitable for larger datasets. In contrast, IVF and PQ indexes offer

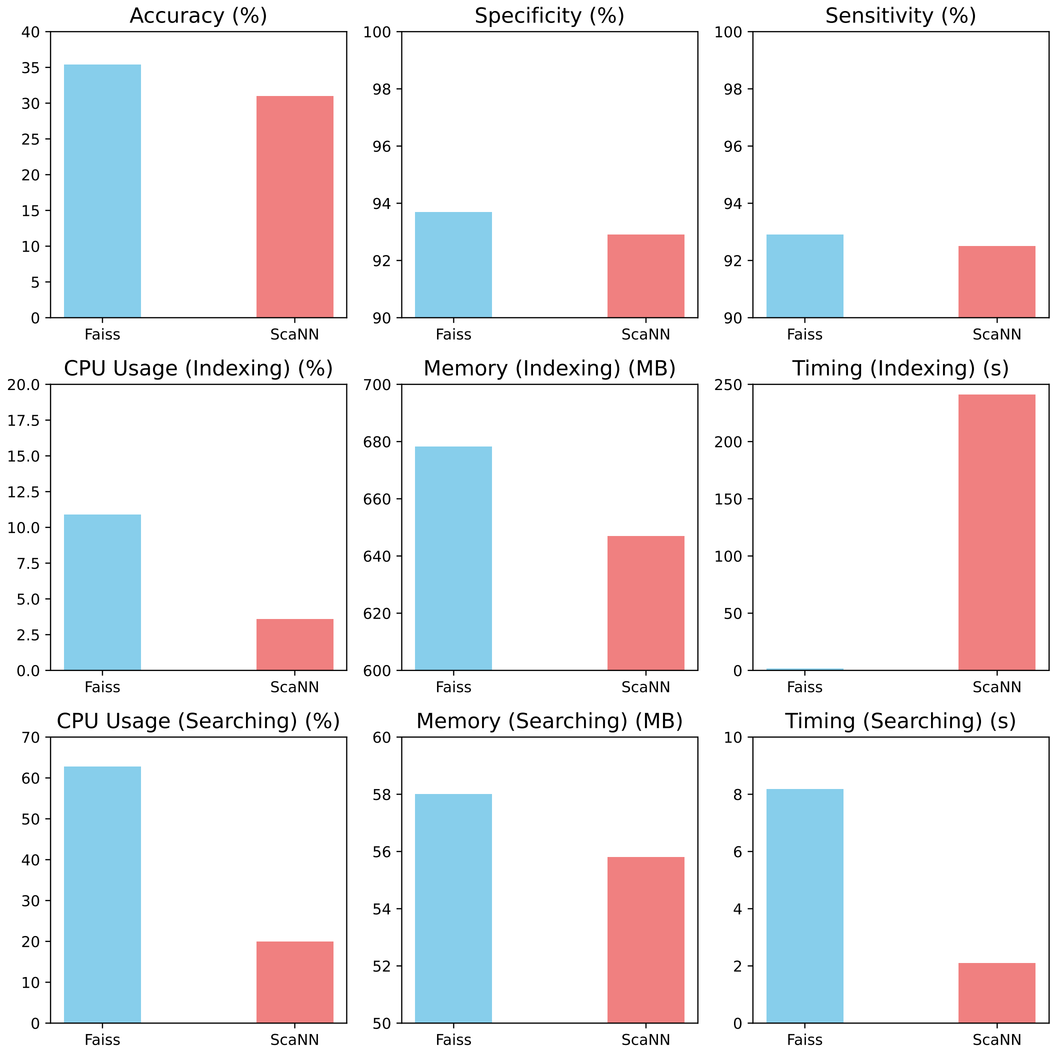


**Figure 12.** Performance Metrics Comparison between Faiss and SCANN.

slower performance and reduced accuracy but are more memory-efficient during indexing. Given the balance between accuracy and general performance, we determined that the most effective indexing method is IVF with a Flat quantizer ("IVF4096, Flat"). However, because our current paper prioritizes accuracy over speed, we have chosen to proceed with the Flat index.

| Index | Time Ind. (s) ↓ | Time Srch. (s) ↓ | CPU Ind. (%) ↓ | CPU Srch. (%) ↓ | Acc. Gene (%) ↑ | Sens. Gene (%) ↑ | Spec. Gene (%) ↑ |
|---|---|---|---|---|---|---|---|
| Flat | 1.6 | 8.2 | 10.9 | 62.8 | 35.4 | 92.9 | 93.7 |
| IMI2x10,Flat | 15.0 | 0.8 | 31.0 | 58.2 | 32.0 | 89.1 | 92.5 |
| IMI2x10,PQ16 | 24.4 | 0.3 | 45.4 | 72.9 | 22.9 | 65.1 | 86.3 |
| IMI2x11,Flat | 35.4 | 0.5 | 37.3 | 42.7 | 31.1 | 85.5 | 92.3 |
| IVF16384,Flat | 59.3 | 1.2 | 39.1 | 27.1 | 34.8 | 92.7 | 93.7 |
| IVF4096,Flat | 24.0 | 4.2 | 36.0 | 46.4 | 35.2 | 92.8 | 93.7 |
| IVF4096,PQ16 | 25.6 | 0.6 | 45.7 | 74.2 | 24.4 | 71.4 | 91.8 |
| OPQ16,IMI2x10,PQ16+16 | 86.4 | 0.2 | 39.4 | 23.7 | 11.3 | 42.9 | 87.2 |
| OPQ32,IVF4096,PQ16x4fsr | 89.0 | 0.1 | 40.8 | 49.9 | 20.6 | 71.8 | 88.2 |
| OPQ32,IVF4096,PQ32 | 98.8 | 0.2 | 40.7 | 24.8 | 22.1 | 75.6 | 91.4 |
| PCA64,IMI2x10,Flat | 4.2 | 0.1 | 14.6 | 92.5 | 7.3 | 75.0 | 94.4 |

**Table 4.** Performance Metrics of FAISS Indexing and Searching Methods: This table presents a comprehensive set of performance metrics for various FAISS configurations.
